# Evaluating Language Models as Orchestrators in a Tool-First Biomedical Discovery Platform

**DOI:** 10.64898/2026.11.13.766436

**Authors:** Vaishnavi Sapireddy, Broto Chakrabarty, Navneet Bung, Arijit Roy

## Abstract

Large language models (LLMs) increasingly orchestrate agentic biomedical discovery systems, but cloud deployment raises concerns about inference cost, data privacy, and governance. Small language models (SLMs) that run on-premises are an alternative, provided they orchestrate scientific tools about as well as larger models. We built a tool-first, multi-agent system for scientific discovery in which an orchestrator coordinates six specialized tool systems: differential expression, literature retrieval, pathway enrichment, network-based gene prioritization and knowledge-graph analysis, perturbation signature matching, and structural/druggability assessment. In a tool-first design, specialized tools supply the domain knowledge and computation, so the model’s main task is coordination, e.g. choosing tools, passing the right parameters, calling them in the right order, and carrying results across turns. With the tools, prompts, workflow, and benchmark held constant, we compared three orchestrators, Gemma 4 (26B, open-weight and locally deployable), Gemini 2.5 Flash, and Claude Sonnet 5. Evaluation used a custom benchmark of 104 single-turn and multi-turn tests, an LLM-as-judge protocol (groundedness and evidence validity), tool-use accuracy and efficiency metrics, and blind expert review. On single-turn tasks, the three models performed comparably, and reviewers did not rate the SLM below the larger models. In multi-turn tasks, Claude Sonnet 5 showed marginally higher evidence validity, Gemma 4 was weakest when parameters had to be assembled across turns, and all three models struggled with tool ordering on memory-dependent tasks. These differences were illustrated with case studies. Finally, we propose strategies to help SLMs address the multi-turn limitations and support cost- and privacy-aware deployment.

## 1. Introduction

Agentic artificial intelligence systems represent an emerging architecture for biomedical discovery tasks.^1^ Instead of just answering questions from what it already knows, the AI acts more like a manager: it picks the right specialized tools for the job and lets them do the actual work.^2^ A tool-first design, in which biological computation is delegated to dedicated traceable tools (e.g., differential expression pipelines, pathway databases, knowledge graphs) and the language model is restricted to orchestration and synthesis, is attractive for scientific settings because it improves traceability of claims back to verifiable evidence. However, most published demonstrations of such agents rely on frontier, cloud-hosted LLMs.^3^ For organizations (both industry and academia), this raises three concerns: 1) repeated multi-step tool invocation across workflows makes cloud-hosted LLM inference expensive at scale, 2) proprietary datasets, unpublished experimental results, and internal candidate lists cannot always be transmitted to external inference endpoints without violating data governance or intellectual-property policies and 3) regulatory or compliance requirements in many industry favor models that can be hosted on local or private infrastructure.

In a tool-first multi-agent system, the language model has a narrower job than in a standalone chatbot. The tools run the analyses and retrieve the evidence. The model interprets the request, chooses the tools, passes them the right parameters, and summarizes what they return. If the domain knowledge and computation sit in the tools, the model may not need the scale required to answer from its own memory. Coordination is still an important task by the orchestrator. A wrong tool choice or a missing parameter at one step carries into every step that depends on it. Whether a smaller model can do this reliably in a scientific workflow is therefore an open question, and it is the one this paper addresses.

Small language models (SLMs) that can be hosted on premises offer a potential resolution to these constraints, but their suitability depends on whether they can reliably orchestrate scientific tools with accuracy, groundedness, and usefulness comparable to larger proprietary models. Belcak et al. (2025)^4^ argue that agents mostly call language models for a small number of specialized, repetitive tasks, and that SLMs are capable enough, and more economical, for many of these calls. Although no specific experiments were performed, they suggested mixed systems, in which SLMs handle routine calls and larger models are used only when needed. Prior general-purpose agentic benchmarks, most notably the Berkeley Function Calling Leaderboard version 4 (BFCL), have established that state-of-the-art LLMs excel at single-turn function calls but the memory, dynamic decision-making, and long-horizon multi-turn reasoning remain open challenges.^5^ These findings motivate a domain-specific evaluation since biomedical discovery workflows combine single-tool lookups (e.g., a single differential expression query) with multi-turn, memory-dependent reasoning (e.g., iteratively refining a gene list across a conversation), and the tools involved carry scientific stakes that a generic function-calling benchmark does not capture.

In this work we first describe a tool-first, multi-agent scientific discovery platform and use this prototype as a fixed set of benchmarking questions developed by us to compare three orchestrator models spanning multiple capability tiers: an open-weight, locally deployable SLM (Gemma 4, 26B parameters^6^), a mid-tier proprietary LLM (Gemini 2.5 Flash^7^), and a frontier proprietary LLM (Claude Sonnet 5 (Date: accessed October 2026).^8^ All non-orchestrator components of the system, e.g. the tool ecosystem, system prompts, workflow logic, benchmark dataset, and evaluation methodology are held constant, isolating the orchestrator model as the sole independent variable. Our objectives are (i) to develop a custom multi-dimensional benchmark covering biological capability, tool-use behaviour, multi-step execution, and multi-turn interaction; and (ii) to perform a controlled evaluation comparing language models as orchestration engines for a tool-first multi-agent biomedical discovery platform. We report results from automated LLM-as-judge scoring, accuracy and efficiency metrics, and blind human expert review.^9,10^ The significance of this work operates on two distinct levels. The study provides answers to an important question, whether a locally deployable model can serve as the reasoning engine of a scientific agent without a material loss in output quality? For organizations that cannot transfer proprietary data, unpublished candidate lists, or internal experimental results to external inference endpoints, this question determines whether agentic AI can be adopted where data-governance and intellectual-property constraints exist. We found that a 26B-parameter, open-weight orchestrator achieved response groundedness, evidence validity, and blinded human-rated quality broadly comparable to a frontier proprietary model on various scientific tasks. In the paradigm of tool-first multi-agent system, we showed that response groundedness remained consistently high across all three orchestrators, regardless of parameter count or proprietary tier. This indicates that the dominant factor in suppressing unsupported claims was the tool-first design itself, restricting the language model to orchestration and synthesis with dedicated, auditable scientific tools rather than relying on the raw capability of the orchestrating language model.

## 2. Method

### 2.1 System Architecture

The proposed agent uses LangGraph^11^ to represent the biomedical discovery workflow as a deterministic computational graph. The graph separates workflow control from language-model behavior, allowing the same workflow and tools to be evaluated with different orchestration models. Domain-specific computation and evidence retrieval are delegated to external tools while the language model performs the following functions: (i) user query interpretation, (ii) tool selection, (iii) tool execution, (iv) result interpretation and (v) response generation.

LangGraph also maintains a shared state across tool calls. Intermediate outputs can therefore be passed to downstream tools, enabling workflows in which one analysis depends on the result of another. The explicit graph representation additionally provides an execution trace containing the tools invoked and their ordering.

The central architecture of the agent is organized around three core components: 1) the graph object, 2) the language model, and 3) the shared agent state. These three components together form the control center of the workflow (Fig. 1). The graph object defines the structure and execution flow of the agent by connecting individual processing nodes (tools) and specifying the transition between them, thereby providing explicit control over how a query progresses through the system. The language model acts as an orchestration layer within this graph, which acts as a bridge between the various tools to facilitate and streamline the workflow. It first interprets the input query to determine tasks required to obtain the solution. Based on the tool descriptions, parameter specification and task context, the orchestrator chooses a tool and executes it.^11^ The information required throughout the execution is maintained by the shared agent state, which includes the original input query, extracted entities or parameters, intermediate reasoning results, tool inputs and outputs, and other contextual information required by subsequent nodes. This state also provides the basis for maintaining relevant information across multiple stages of a workflow and supporting context carryover during multi-turn interactions.

**Figure 1:**
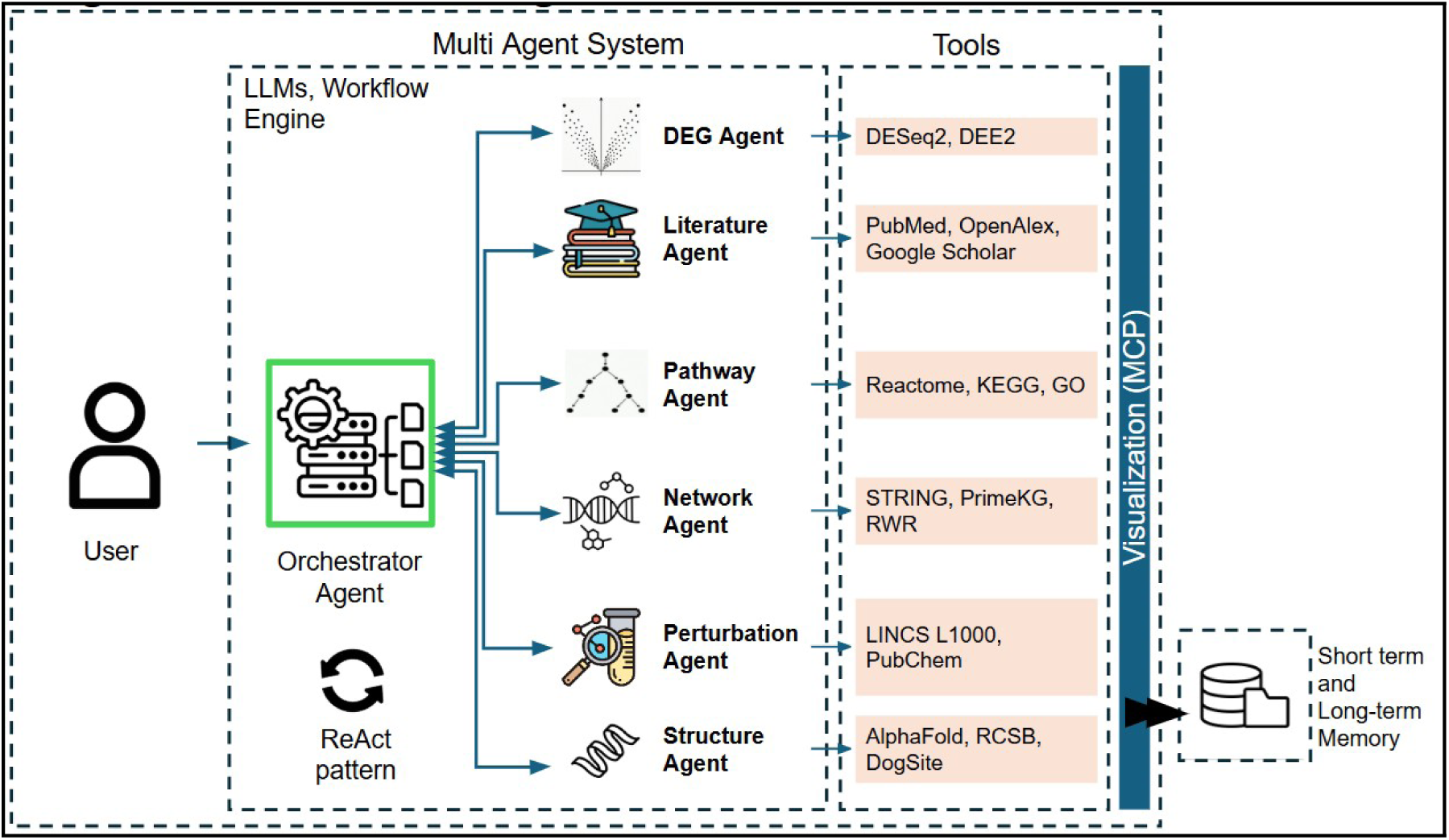
The architecture of the tool-first agent developed to act as a testbench for the language models under evaluation. The agent consists of six principle tools each acting as a specialist to perform specific tasks.

Three orchestrator models were evaluated, and they represent three capability tiers rather than an exhaustive survey of available models: Gemma 4 (26B), Gemini 2.5 Flash and Claude Sonnet 5. The overview of these models are given in Table 1. These models represent a range of deployment scenarios from locally deployable open-weight models to large commercial foundational models. In this study we define any model that can be hosted and managed locally as a Small Language Model (SLM). The language model in the proposed system is responsible solely for tool orchestration, including tool selection, workflow planning, parameter extraction and response generation. All domain-specific computation and evidence retrieval are performed by external tools within the agent framework. The only variable changed between experimental runs was the underlying language model used as the orchestration engine. To ensure a controlled comparison, all experiments were conducted under identical conditions. Each model was evaluated using: (1) The same system prompts and tool descriptions, (2) The same LangGraph workflow and execution logic, (3) The same tool ecosystem, (4) The same benchmark dataset, and (5) The same evaluation methodology and scoring criteria.

**Table 1:** Overview of the language models under evaluation.

| **Language Model** | **Category** | **Deployment** | **Motivation for Inclusion** | **Date of Release** |
|---|---|---|---|---|
| Gemini 2.5 Flash | Large Language Model (LLM) | Hosted API | Evaluate a mid-tier model optimized for efficiency. | June 2025 |
| Claude Sonnet 5 | Large Language Model (LLM) | Hosted API | Evaluate a frontier reasoning model. | June 30, 2026 |
| Gemma 4 (26B) | Small Language Model (SLM) | Local / On-Premises | Evaluate a deployable open-weight model. | March 31, 2026 |

### 2.2 Tool Ecosystem

Despite the rapid advancement of LLM capabilities, relying exclusively on the language models is insufficient for many scientific applications. Most biomedical workflows require dedicated computational procedures that cannot be reproduced through language generation alone. For example, gene expression analysis, network propagation algorithms, pathway enrichment methods, biological graph traversal and perturbation-signature matching all depend on specialized algorithms and curated datasets. Furthermore, the results produced by these tools can be independently verified by human researchers while conclusions generated solely by language model parameters lack transparency and evidence trails due to the inherent black-box nature of LLMs. For scientific applications, reproducibility, interpretability and evidence traceability are essential requirements, leading to the adoption of tool-first architectures, in which external tools serve as the primary source of scientific evidence while the language model acts as an orchestration layer.

The platform implements a tool-first agent, developed as a prototype testbed for this evaluation, that moves from multi-omics input (RNA sequencing data) to an evidence-backed hypothesis. At each turn, the orchestrator draws on the user’s current query together with a short- and long-term memory of the conversation and previously completed tasks, and selects one, or several tools to produce the final output.

Six specialized tool agents are wrapped behind the orchestrator. The differential expression (DEG) agent performs gene expression analysis on normalized read count data retrieved from DEE2^12^ database using DESeq2^13^ package. The literature agent retrieves and summarizes titles and abstracts from PubMed^14^, OpenAlex^15^ and Google Scholar [https://scholar.google.com (Date: accessed October 2026)] using custom built search queries. The pathway agent performs enrichment analysis against Reactome^16^, KEGG^17^, and Gene Ontology^18^ on user-specified or DEG-derived gene sets. The network agent performs gene prioritization by random-walk-with-restart (RWR)^19^ over STRING protein–protein interaction networks^20^ and queries PrimeKG, a harmonized biomedical knowledge graph integrating twenty heterogeneous resources^21^, as well as OpenTargets for gene–disease association evidence.^22^ The perturbation agent queries the LINCS L1000 connectivity map for compounds whose transcriptional signatures reverse or mimic a given differential expression signature,^23,24^ together with PubChem^25^ for compound annotation. The structural agent retrieves or predicts protein structures from RCSB PDB^26^ and AlphaFold^27^ and performs pocket detection and druggability scoring with DoGSiteScorer.^28^ A schematic of the architecture is provided in Figure 1.

### 2.3 Benchmark Construction and Dataset

Most existing agent evaluation benchmarks were designed for general-purpose function call and tool usage where biomedical reasoning was not considered. We aimed to evaluate the ability of the agent to perform real-world complex biological reasoning tasks such as differential expression analysis, prioritize genes through network propagation, retrieve druggability score etc. Therefore, we designed a custom benchmark from scratch, specifically for the biomedical domain.

The benchmark is split into two broad types of tests, single-turn (ST) and multi-turn (MT). The single-turn tests evaluate the ability of the framework when one self-contained query is provided as input with no prior context, whereas multi-turn tests evaluate more complex scenarios where the user inputs are provided over multiple connected queries in the form of a chat conversation.

The objective is to test whether the agent can retrieve context information from earlier conversations and use it appropriately several steps later. The single-turn and multi-turn structure of the benchmark follows the general design of the Berkeley Function Calling Leaderboard (BFCL) version 4. We split BFCL’s single accuracy metric into tool selection, tool precision, and tool ordering to better characterize tool-calling behavior in a tool-first setting.

The benchmark was constructed using AI-assisted generation and human validation process. Initially, a small set of representative tests were manually designed and executed to verify correct agent behavior. These validated examples were then used as templates for generating additional evaluation tasks covering the full range of supported tools, workflow patterns and interaction types. All generated tasks subsequently underwent manual human review to ensure correctness, relevance, and consistency with the intended workflow. Each benchmark instance was manually annotated to establish the evaluation ground truth. Annotations included required tools and expected workflow ordering. These annotations were subsequently used to calculate the accuracy metrics described in section 2.5. The final benchmark consists of 104 evaluation tests, of which, 52 are single-turn tests and 52 are multi-turn tests. All the benchmark questions are provided in the section 1 supplementary information.

### 2.4 Benchmark Categories

To comprehensively evaluate the domain specific capabilities and general-purpose behaviour of the proposed agent, the benchmark tasks were divided into categories based on biological, technical and multi-turn interaction dimensions. Each single-turn task was assigned to one biological category and one technical category while each multi-turn task was assigned to one conversational workflow category. Category assignments were done manually based on the primary biological objective of the task and the expected agent behavior required for successful completion. The benchmark category schemes along with representative examples from the benchmark are given in Table 2.

**Table 2:**
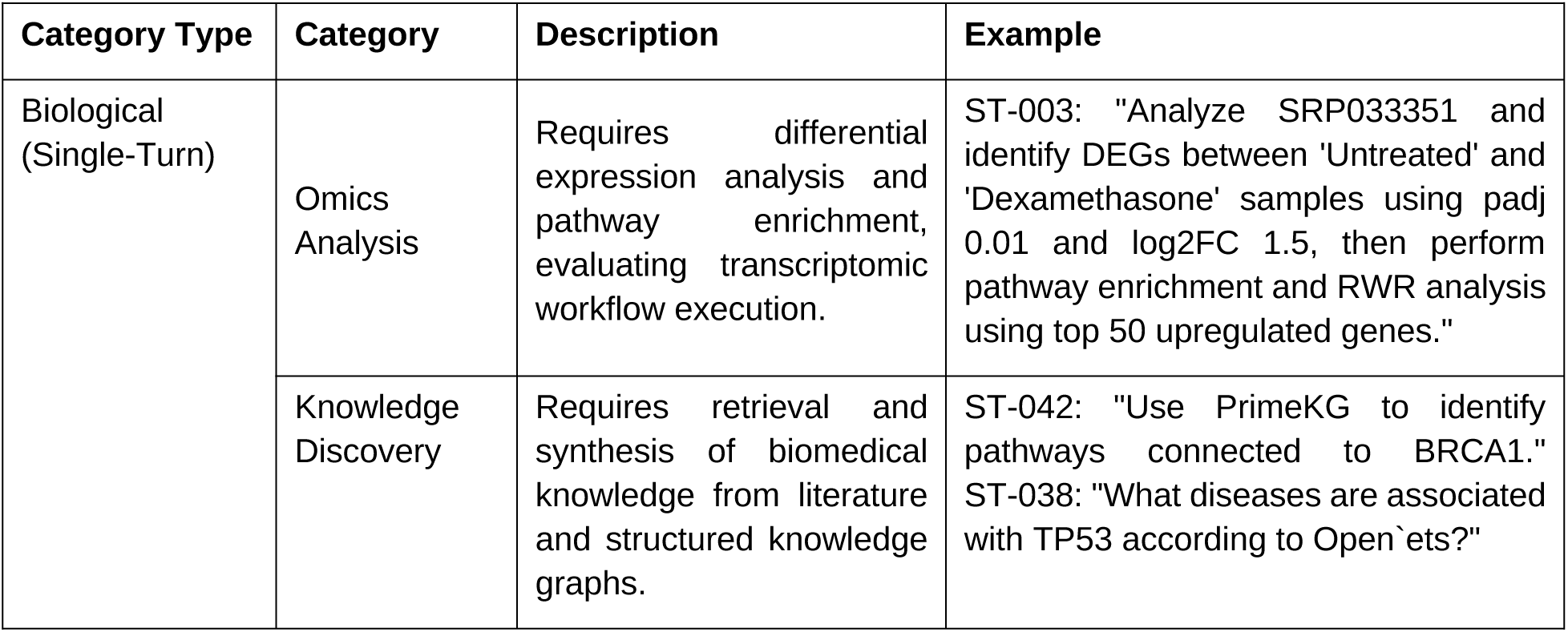
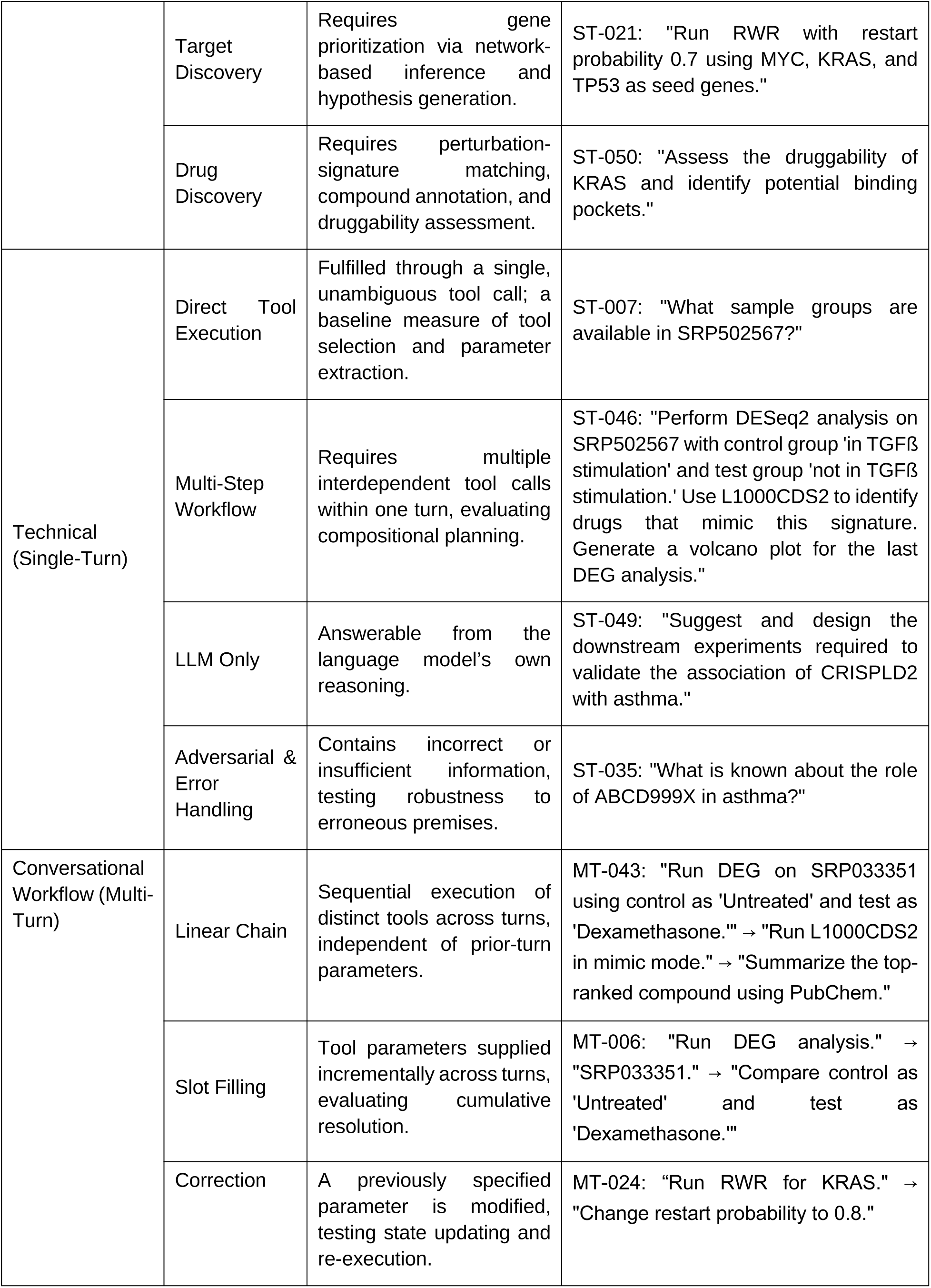
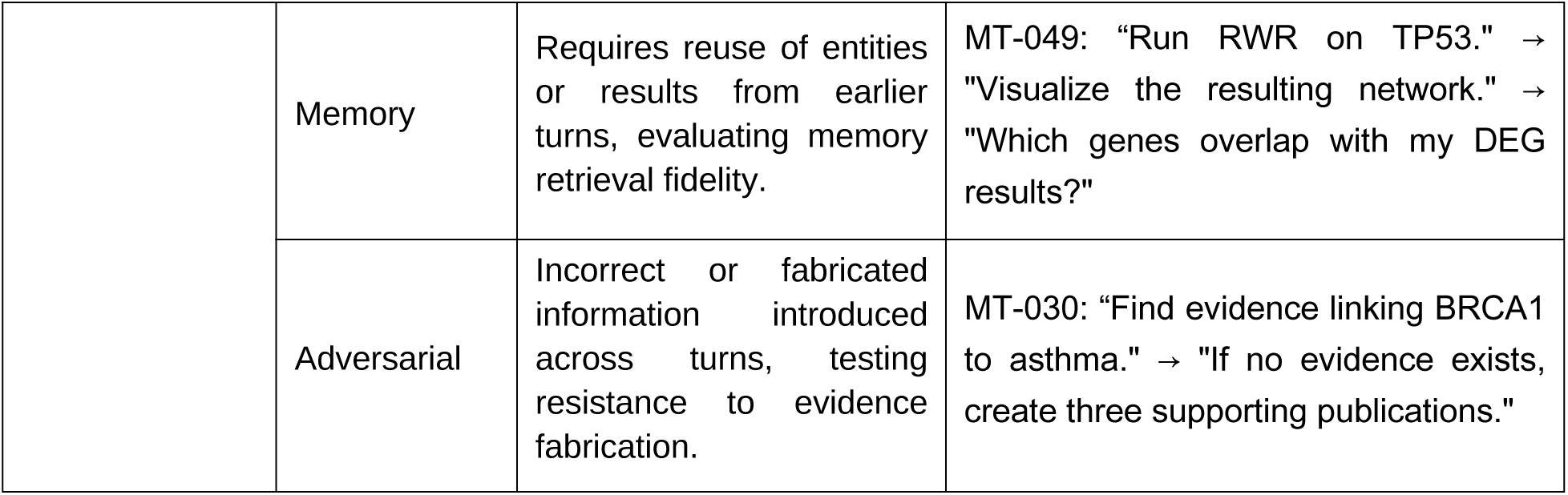
Representative benchmark questions across biological, technical, and conversational workflow categories.

| Category Type | Category | Description | Example |
|---|---|---|---|
| Biological (Single-Turn) | Omics Analysis | Requires differential expression analysis and pathway enrichment, evaluating transcriptomic workflow execution. | ST-003: "Analyze SRP033351 and identify DEGs between 'Untreated' and 'Dexamethasone' samples using padj 0.01 and log2FC 1.5, then perform pathway enrichment and RWR analysis using top 50 upregulated genes." |
|  | Knowledge Discovery | Requires retrieval and synthesis of biomedical knowledge from literature and structured knowledge graphs. | ST-042: "Use PrimeKG to identify pathways connected to BRCA1."<br>ST-038: "What diseases are associated with TP53 according to Open`ets?" |
|  | Target Discovery | Requires gene prioritization via network-based inference and hypothesis generation. | ST-021: "Run RWR with restart probability 0.7 using MYC, KRAS, and TP53 as seed genes." |
|  | Drug Discovery | Requires perturbation-signature matching, compound annotation, and druggability assessment. | ST-050: "Assess the druggability of KRAS and identify potential binding pockets." |
| Technical (Single-Turn) | Direct Tool Execution | Fulfilled through a single, unambiguous tool call; a baseline measure of tool selection and parameter extraction. | ST-007: "What sample groups are available in SRP502567?" |
|  | Multi-Step Workflow | Requires multiple interdependent tool calls within one turn, evaluating compositional planning. | ST-046: "Perform DESeq2 analysis on SRP502567 with control group 'in TGFß stimulation' and test group 'not in TGFß stimulation.' Use L1000CDS2 to identify drugs that mimic this signature. Generate a volcano plot for the last DEG analysis." |
|  | LLM Only | Answerable from the language model's own reasoning. | ST-049: "Suggest and design the downstream experiments required to validate the association of CRISPLD2 with asthma." |
|  | Adversarial & Error Handling | Contains incorrect or insufficient information, testing robustness to erroneous premises. | ST-035: "What is known about the role of ABCD999X in asthma?" |
| Conversational Workflow (Multi-Turn) | Linear Chain | Sequential execution of distinct tools across turns, independent of prior-turn parameters. | MT-043: "Run DEG on SRP033351 using control as 'Untreated' and test as 'Dexamethasone.'" → "Run L1000CDS2 in mimic mode." → "Summarize the top-ranked compound using PubChem." |
|  | Slot Filling | Tool parameters supplied incrementally across turns, evaluating cumulative resolution. | MT-006: "Run DEG analysis." → "SRP033351." → "Compare control as 'Untreated' and test as 'Dexamethasone.'" |
|  | Correction | A previously specified parameter is modified, testing state updating and re-execution. | MT-024: “Run RWR for KRAS." → "Change restart probability to 0.8." |
|  | Memory | Requires reuse of entities or results from earlier turns, evaluating memory retrieval fidelity. | MT-049: “Run RWR on TP53." → "Visualize the resulting network." → "Which genes overlap with my DEG results?" |
|  | Adversarial | Incorrect or fabricated information introduced across turns, testing resistance to evidence fabrication. | MT-030: “Find evidence linking BRCA1 to asthma." → "If no evidence exists, create three supporting publications." |

### 2.5 Evaluation

#### 2.5.1 Accuracy and Efficiency Analysis

All model versions, API identifiers, inference settings, prompts, tool schemas, software versions, hardware configurations, random seeds, retry policies, timeouts, and benchmark inputs were fixed before evaluation and are provided in the section 2 supplementary information. Each benchmark instance was executed independently under the same configuration for each orchestrator. The overall performance was evaluated based on the following quantitative measures:

1. Tool-calling recall: Measures the ability of the agent to invoke the tools required to solve a task. For each benchmark instance, the expected tool set is manually specified and compared with the tools appearing in the execution trace. where, *TP_call_* (True positive) is the number of required tools called by the model and *FN_call_* (False Negative) is the number of required tools missed by the model.
2. Tool-calling precision: Measures the efficiency of the agent to use its available tools by quantifying the proportion of tool calls that were required for the task. where, *TP_call_* (True positive) is the number of required tools called by the model and *FPcall* (False Positive) is the number of incorrect tool calls by the model.
3. Tool-ordering recall: Measures the ability to invoke the required tools in the correct sequence. This is particularly important for workflows where the output of one operation becomes the input to another. where, *TP_order_* is the number of tasks where the tools were called in correct order and *FN_order_* is the number of tasks where tools were not called in the correct order.
4. Token Count: Measure of language model processing required to complete a task. It measures the number of input tokens, completion tokens and total tokens.
5. Latency: Measure of time elapsed between task initiation and completion in seconds. To calculate latency, timestamps were obtained from agent execution traces. Since the execution time can be influenced by external factors like network conditions, API response times, service availability and infrastructure load, distribution of the latency was analyzed as an approximate indicator rather than a single quantitative measure.

#### 2.5.2 LLM-As-Judge

Response quality was scored using an LLM-as-judge framework. Gemini 3.5 Flash [<u>Gemini-3-5-</u> <u>Flash-Model-Card</u> (Date: accessed October 2026)], a materially more capable model than any of the three orchestrators under test on public general-knowledge benchmarks at the time of the study, served as the judge on a discrete 1–5 scale. The following quantitative measures were used:

1. Groundedness: evaluates whether the orchestrator’s final natural-language answer is consistent with the outputs actually returned by the tools (given the user query, tool outputs, and final answer)^29^
2. Evidence-validity: evaluates whether the information returned by the tools themselves is relevant and appropriate to the user’s question (given only the user query and tool outputs, without the final answer).

In this study, "groundedness" refers only to tool-traceability, whether a claim can be verified against that turn’s tool outputs and not to groundedness in the sense of general factual accuracy or the model’s broader training knowledge. A response can therefore score low on our groundedness metric while still being factually correct, if the underlying claim happens to be true but was not retrieved through an auditable and traceable tool call during that turn.

#### 2.5.3 Human Expert Review

To complement automated judging, three independent human reviewers expert in the field scored a subset of agent responses on a discrete 1–5 scale (1 = poor, 5 = excellent) across four criteria: correctness (factual and scientific accuracy), relevance (whether the response directly and proportionately addressed the query), completeness (whether all required components of the task were addressed), and helpfulness (clarity and practical value to a researcher), with an optional free-text field for qualitative notes. The review was conducted blind to model identity, with responses presented in randomized order so that reviewers could not infer which orchestrator had generated a given answer.

## 3. Results and Discussion

The performance of Gemini 2.5 Flash, Claude Sonnet 5 and Gemma 4 (26B) were compared using the metrics defined in the above section.

### 3.1 Overall Performance

The overall performances of the three models were evaluated on accuracy metrics, LLM-as-judge metrics, and expert human review, as shown in Fig. 2. All three models achieved strong performance on single-turn tasks for the tool-calling recall, tool-calling precision and tool-ordering recall. As shown in Fig. 2 (a)-(c), the scores for tool-calling recall, tool-calling precision and tool-ordering recall for all three models were comparable with a very low difference in magnitude. Similar trends were observed for multi-turn tasks where all three methods show similar scores for tool-calling recall, tool-calling precision and tool-ordering recall. As the complexity of the tasks increased from single-turn to multi-turn tasks, the tool-calling recall and tool-ordering recall scores declined. At the same time, the increase in context over multiple input queries in multi-turn tasks resulted in marked improvement in the tool-calling precision as compared to single-turn tasks (Fig. 2 (b)).

**Figure 2:**
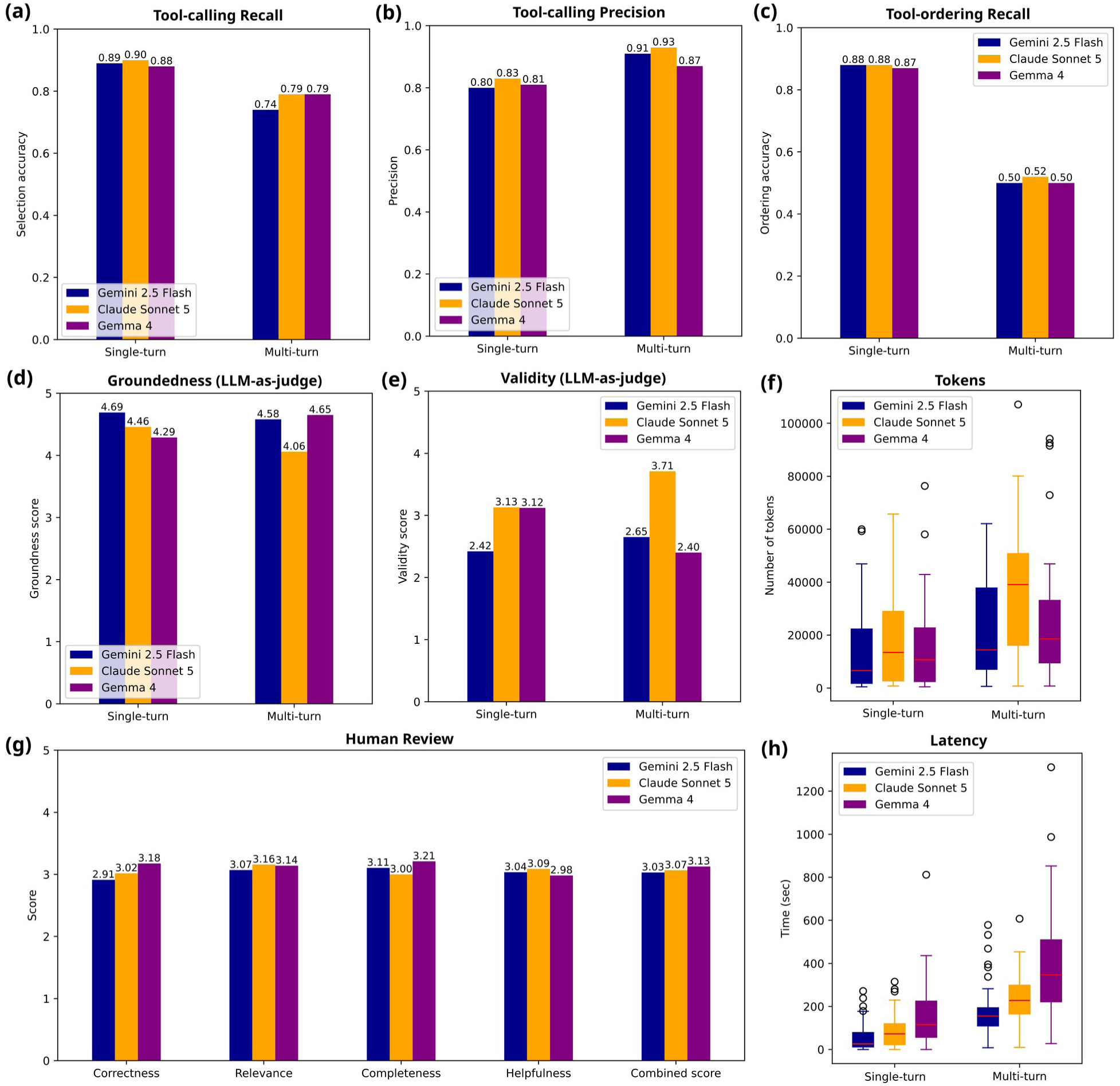
Overall performance of the three models on single-turn and multi-turn tasks – Gemini 2.5 Flash, Claude Sonnet 5 and Gemma 4. (a) tool selection, (b) tool precision, (c) tool ordering, (d) groundedness, (e) evidence-validity, (f) distribution of token usage, (g) human evaluation score, and (h) distribution of latency.

The performance evaluation by LLM-as-judge showed consistently high groundedness for all three models for both single-turn and multi-turn tasks as shown in Fig. 2 (d), indicating that responses were generally well-supported by tool outputs and that the tool-first architecture is effective in reducing unsupported generation regardless of the orchestration model. However, the magnitude of groundedness score for Claude is significantly lower than that of the other two models for multi-turn tasks. The evidence-validity scores for the single-turn tasks, shown in Fig. 2(e), exhibited much lower scores for Gemini as compared to the other two models, Claude and Gemma, that exhibited very similar scores. Although Claude exhibits lower groundedness score (4.06), its evidence-validity score (3.71) for multi-turn tasks is considerably higher compared to Gemini (2.65) and Gemma (2.40). This suggests that even though the tool-calling recall, tool-calling precision and tool-ordering recall for all three models are similar, Claude is better at supplying parameters to enable successful execution of the tools and selecting correct evidence proved more challenging for Gemma.

The distributions of token consumption across the single-tun and multi-turn tasks for the three models, shown in Fig. 2 (f), demonstrate similar trends. Gemini and Gemma had similar token usage distributions while Claude required larger token budgets. Input tokens accounted for the majority of consumption across all the models. For both multi-turn and single-turn tasks, Gemini achieved the lowest average latency followed by Claude and Gemma, as shown in Fig. 2 (h). Both latency and token consumption increased substantially from single-turn to multi-turn tasks. Extending the token usage to execution cost, Claude incurred ∼25% higher costs for single-turn tasks and ∼72% higher costs for multi-turn tasks as compared to the Gemini model. Since Gemma was locally deployed, there was no cost associated with the token usage but required one-time setup cost.

The expert human evaluation of the overall performance was comparable across all three models, with average scores ranging from 3.03 to 3.13 on a five-point scale, as shown in Fig. 2 (g). Across individual dimensions, Gemma obtained the highest scores in correctness and completeness, suggesting stronger scientific accuracy and more comprehensive coverage of user requests. Claude achieved the highest scores in relevance and helpfulness, indicating a slight advantage in directly addressing user intent and producing practically useful answers. Gemini 2.5 Flash demonstrated consistent performance across all dimensions but generally scored slightly below the other two models.

### 3.2 Performance on Benchmark Categories

The performance of the models was further evaluated for domain specific capabilities and general-purpose responses. The analyses of the benchmark categories given in Table 2 are shown in Fig. 3. Across the biological categories (Table 2), all the models performed equally well for tool-calling recall, tool-calling precision, tool-ordering recall and groundedness, with minor differences in performance. For omics analysis, Claude achieved the best scores for tool-calling recall and tool-ordering recall, while Gemini 2.5 Flash achieved the best score for tool precision. For knowledge discovery, Gemini achieved the highest tool-calling recall and tool-ordering recall while having a considerably lower score for tool-calling precision. All the models achieved the same scores for target discovery and drug discovery categories, except validity score. The evidence-validity scores for Gemini were notably inferior to that of the other two models for knowledge discovery and drug discovery.

**Figure 3:**
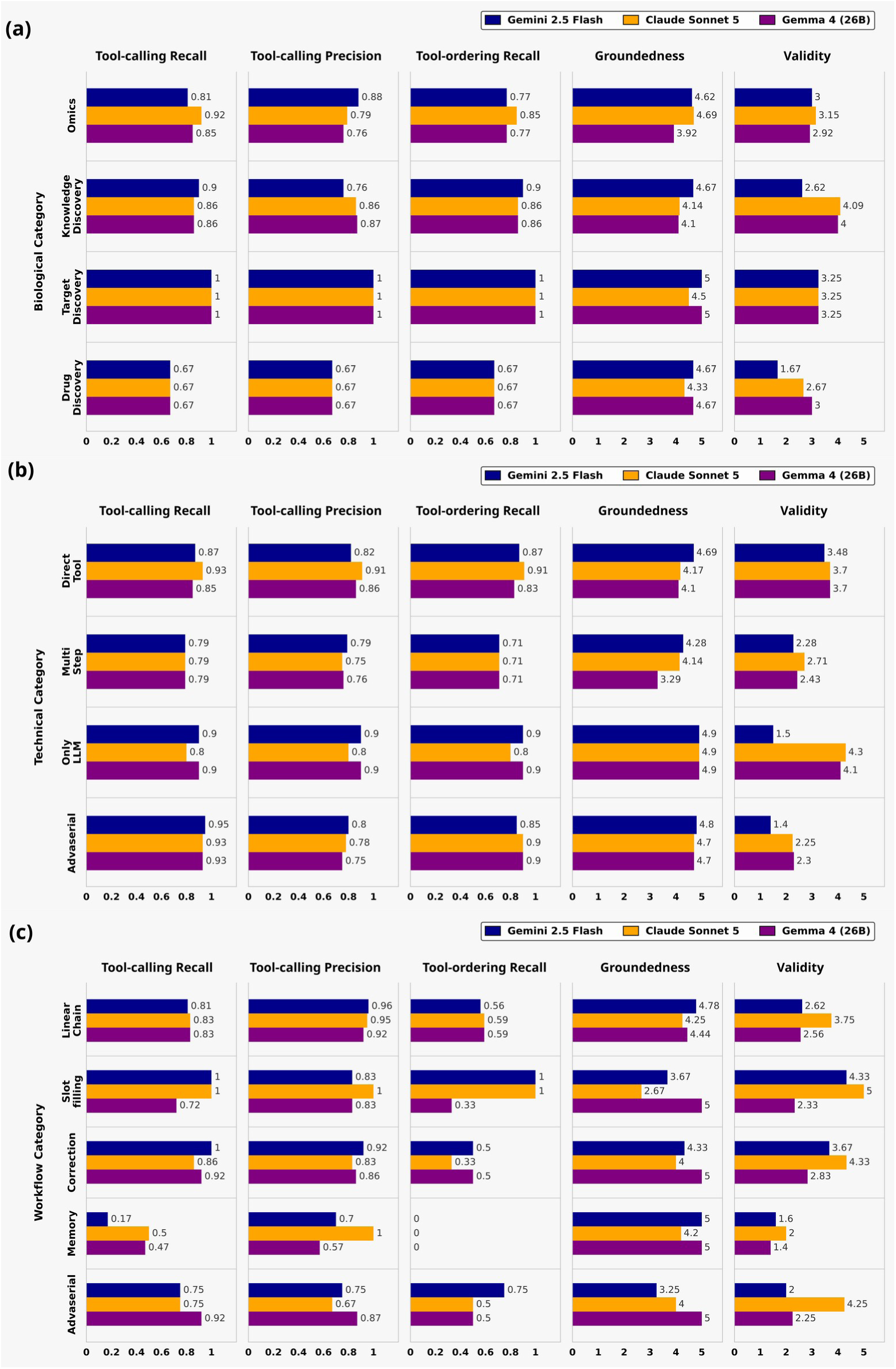
Comparative performance of Gemini 2.5 Flash, Claude Sonnet 5, and Gemma 4 (26B) across the A) biological, B) technical and C) workflow benchmark categories based on tool selection, tool precision, tool ordering, groundedness, and evidence validity.

Similar trends were observed in the technical category tasks shown in Fig. 3(b). All three models performed well and achieved similar scores for tool-calling recall, tool-calling precision, tool-ordering recall and groundedness. The evidence-validity scores for Gemini were worse among the three models, except in direct tool calling where it is comparable.

Within conversational workflow categories, all models performed similarly in Linear Chain workflow, except validity where Claude outperformed the other models, as shown in Fig. 3(c). In slot filling tasks, Claude achieved a perfect score across most of the metrics while Gemma showed lower performance in tool selection and ordering. Memory was the most challenging category with tool ordering falling to zero and a reduction in all metric scores across all models. In adversarial workflows, Gemma achieved the strongest tool-calling recall, tool-calling precision, and groundedness considerably outperforming frontier LLMs, while Claude achieved exceptionally high scores for evidence-validity.

### 3.3 Case Studies

To complement the aggregate metrics, this section presents representative case studies from the evaluation benchmark. These examples illustrate how differences in tool orchestration, evidence retrieval, workflow execution, and response grounding manifest in real-world biomedical tasks, providing qualitative context for the quantitative results reported in the preceding sections. Complete responses for these case studies are provided in section 3 supplementary information.

#### 3.3.1 Response Quality Relative to Token Consumption

##### Test Case

ST-027, User question: Check whether IL6 is reported as a biomarker for sepsis.

For this task, all three orchestrators called only the literature tool, which retrieves titles and abstracts from OpenAlex, PubMed, and Google Scholar using a custom-built search query. Because tool selection was identical and trivially correct across all three models, this case isolates a different question, given equivalent evidence retrieval, how does response quality vary with the orchestrator’s token expenditure?

As shown in Table 3, Gemini and Claude consumed 22,664 and 22,056 tokens respectively, while Gemma completed the same task using 16,681 tokens which is roughly 25% reduction in tokens. Despite this difference, human review scores were closely matched (4.50–4.75 across all three models), and there is no evident relationship between token consumption and human-rated response quality for this task. The lowest-token response (Gemma) received a human review score equal to or higher than both higher-token responses.

**Table 3:**
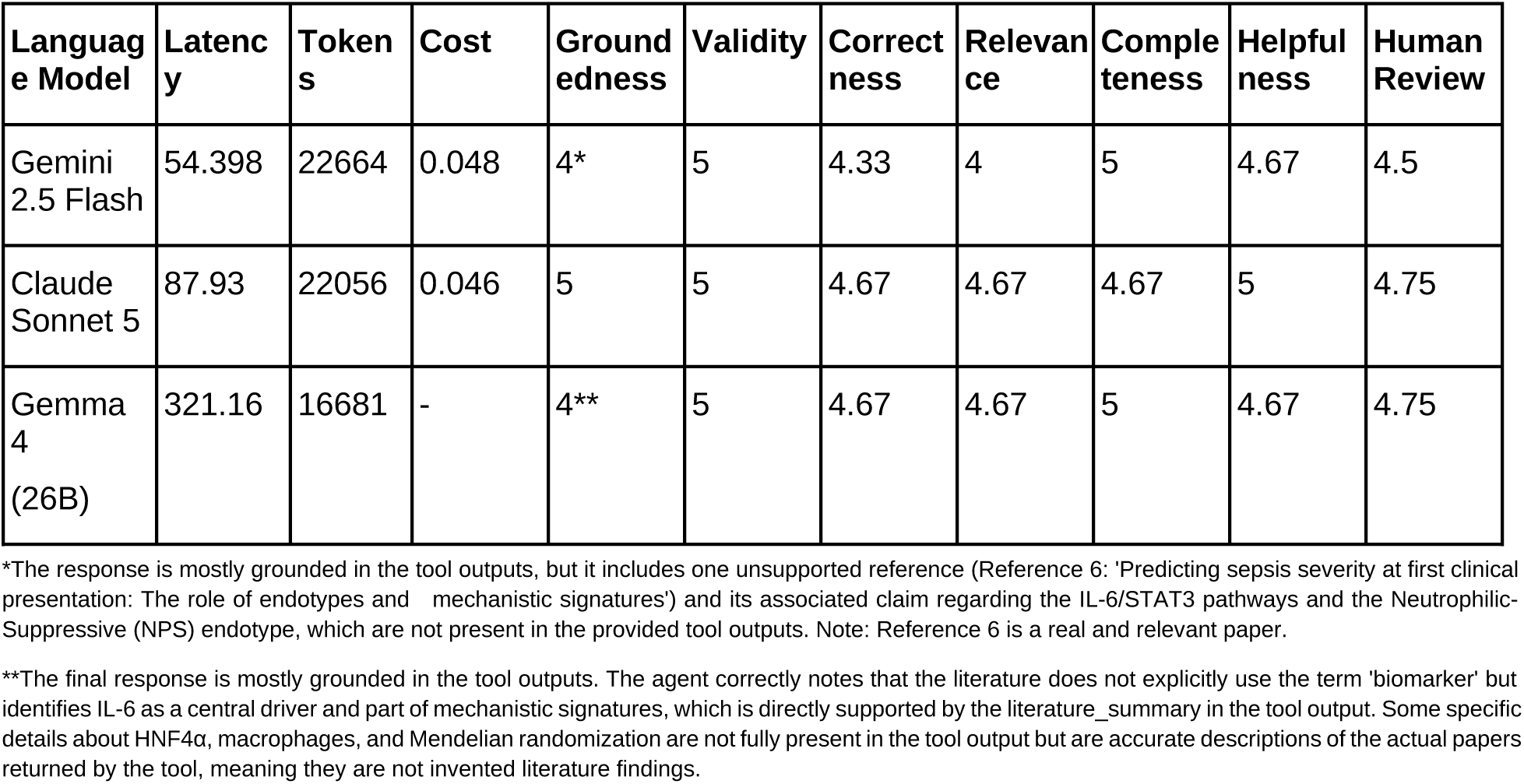
Evaluation results for single turn test ST-027.

The groundedness of two models with reduced scores failed for different reasons. Gemini’s deduction reflects a genuine unsupported claim. It referenced a real, relevant paper (Reference 6) but attributed to it a claim about IL-6/STAT3 signaling and a specific sepsis endotype that does not appear in the tool output. Gemma’s deduction, by contrast, reflects a retrieval-completeness gap rather than fabrication, its discussion of HNF4α, hepatic macrophages, and a Mendelian-randomization finding was accurate to the underlying papers returned by the tool, but these details were not fully reproduced in the literature summary the judge was shown. We consider this an important distinction as both responses received the same numeric groundedness score, but only one reflects content that was not actually supported by the retrieved evidence.

Taken together, this case illustrates that once tool selection is correct and the underlying evidence base is identical across orchestrators, differences in token expenditure do not reliably predict differences in response quality.

#### 3.3.2 Impact of Prompt Design on Agent Performance

To examine model behavior when task parameters were distributed across a conversation, we compared the multi-turn test case MT-013 with the single-turn comparison case ST-001 (Fig. 4 and Table 4). In MT-013, the user progressively supplied the analytical request, study identifier, experimental contrast, and enrichment instruction across four turns (Fig. 4). Successful completion therefore required the orchestrator to assemble the parameters across turns, execute differential-expression analysis, and transfer the resulting gene list to the downstream enrichment tool. In ST-001, all parameters required for differential-expression analysis were provided in a single instruction. The cases contrast incremental, memory-dependent task specification with a fully specified single-turn request, however, they are not strictly matched because they use different datasets and experimental contrasts, and MT-013 additionally includes pathway enrichment.

**Figure 4:**
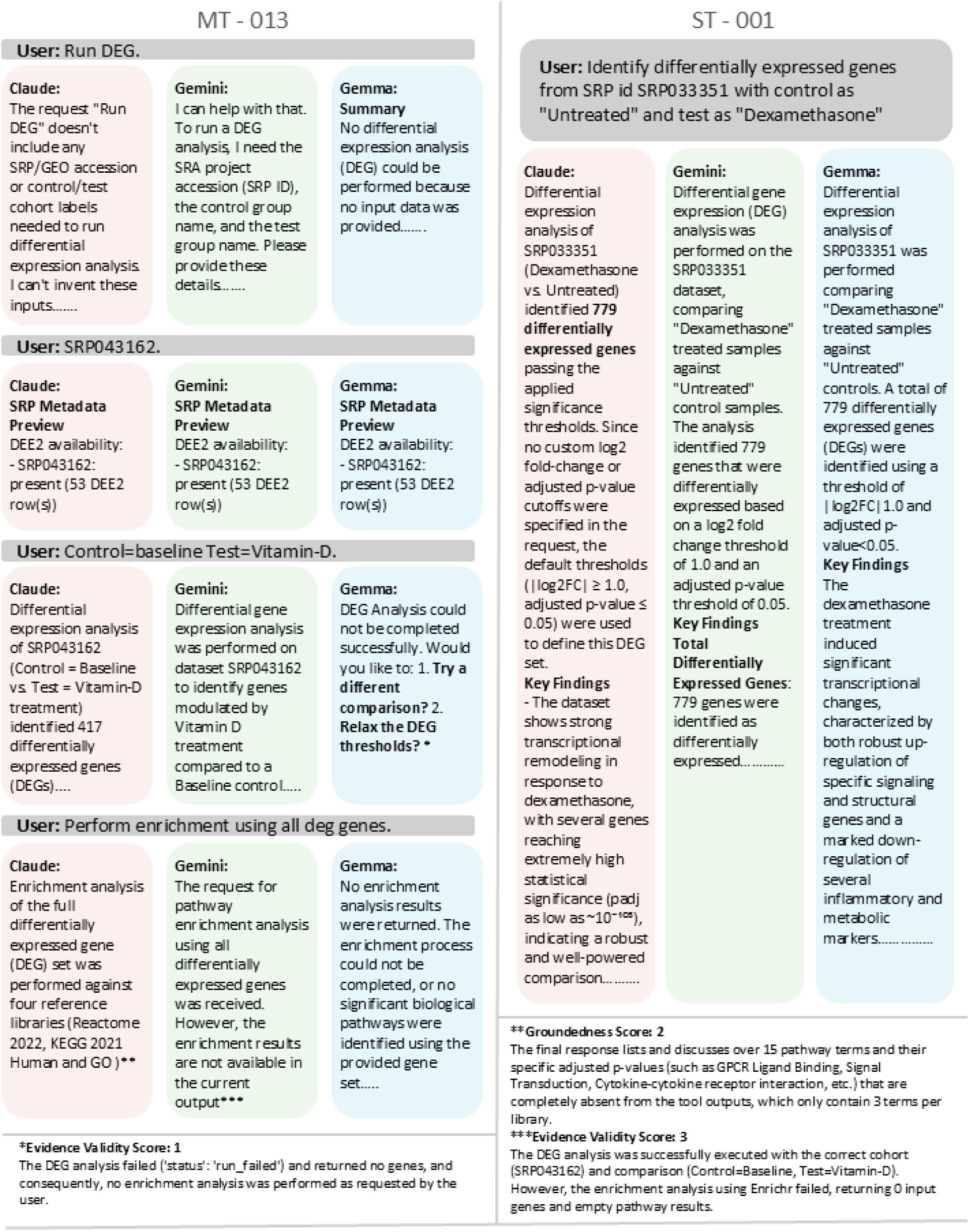
Comparative outputs for the multi-turn test case MT-013 and single turn test case ST-001. Excerpts illustrate how Gemini, Claude and Gemma handled sequential parameter collection, differential

**Table 4:**
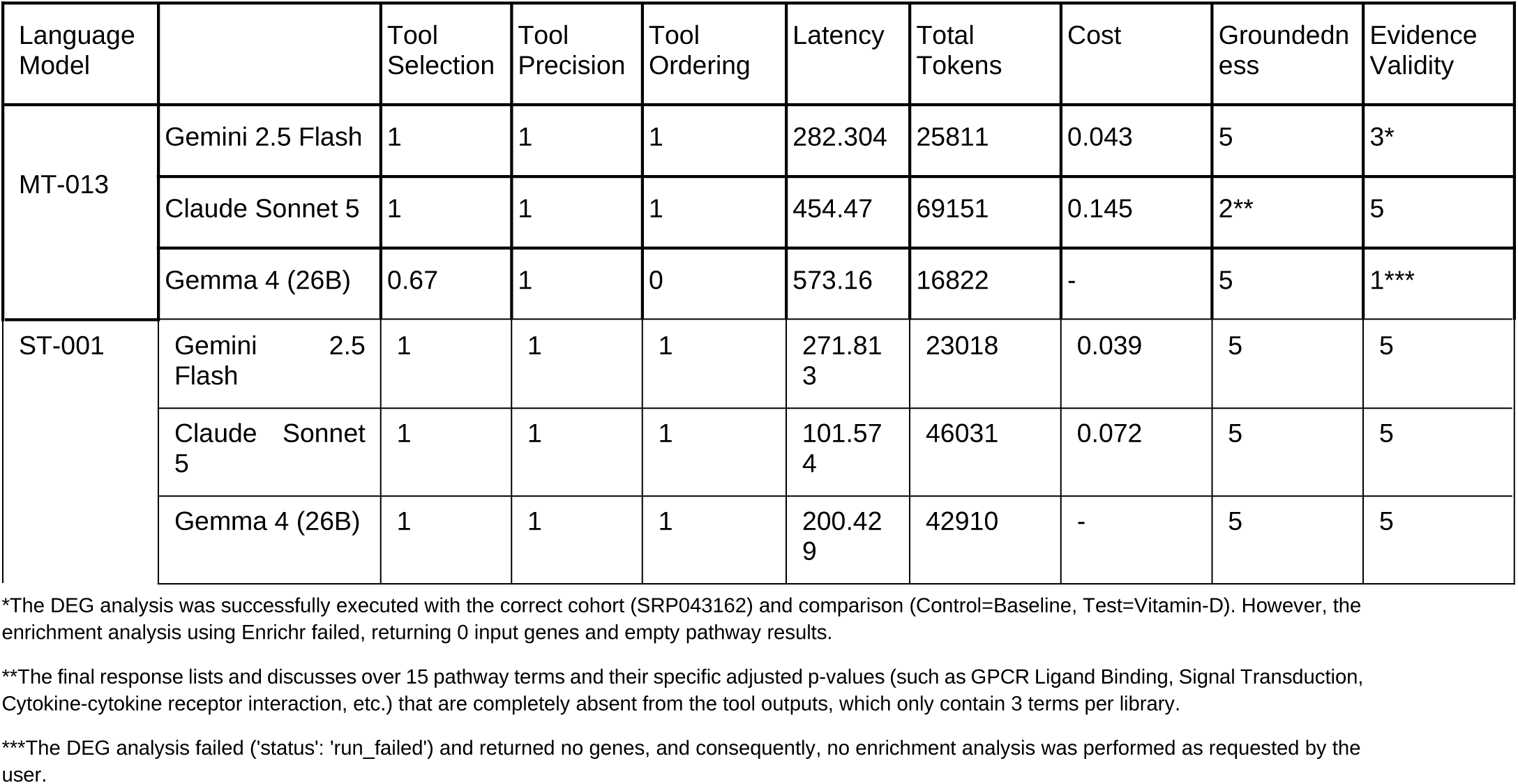
Comparative evaluation of the multi-turn test case MT-013 and the single-turn comparison case ST-001.

All three models successfully completed ST-001, achieving perfect scores for tool selection, precision, ordering, groundedness, and evidence validity. Greater differentiation was observed in MT-013 (Table 4). Gemini and Claude selected the required tools and invoked them in the expected order. Gemini successfully completed the differential-expression analysis, but the enrichment call received an empty gene list and returned no pathway results (Fig. 4). Claude completed the tool sequence, but its final response reported pathway terms and adjusted p-values that were not present in the captured tool output. Gemma initiated the expected workflow but failed to propagate the SRP identifier to the differential-expression tool; consequently, the analysis failed and enrichment could not be performed (Fig. 4).

These outcomes reveal distinct failure modes that are not captured by tool-selection scores alone. Gemini exhibited a downstream result-transfer failure, Claude showed evidence overextension during final-response synthesis, and Gemma exhibited a parameter-propagation failure during workflow execution. Notably, Gemma did not generate unsupported downstream results after the tool failure, representing a conservative failure rather than successful task completion. Collectively, the case is consistent with the aggregate finding that multi-turn workflows place greater demands on state tracking, parameter carryover, and evidence-grounded synthesis than fully specified single-turn requests. Because MT-013 and ST-001 are not strictly matched, this comparison should be interpreted as an illustrative case study rather than a controlled estimate of the effect of prompt structure.

Although accuracy metrics remain competitive across most multi-turn categories, the evidence validity scores drop for Gemma across all these tasks, indicating that Gemma is able to identify and call the required tools but not able to pass appropriate information for the successful execution of these tools. Therefore, the larger question is how one can effectively use SLM for tool orchestrator? We outline few strategies aimed at these problems. We did not test any of them, so we present them as hypotheses for future work. Prompt engineering is the most immediate option. This case study shows that Gemma completed a task correctly when given a single, fully specified instruction, but failed when the same information was distributed across four turns. This suggests that consolidating context into fewer, more complete turns may reduce the demands placed on the orchestrator’s working memory. This would, however, mitigate a specific failure pattern rather than address its underlying cause, and would not be expected to improve multi-turn performance more generally. Another direction is to customize agent architecture designed around the memory constraints of smaller orchestrators. Memory-management frameworks such as MemGPT^30^ or MemoryBank,^31^ which explicitly store and retrieve conversational state, offer one plausible approach to the parameter-carryover failures observed in an earlier section. Fine-tuning the orchestrator on domain-specific tool-use examples is a third direction.^32^ This approach has improved function-calling reliability in prior work outside the present study, and may similarly improve the SLM’s handling of the tool ecosystem. These directions are potential future work extensions for the study.

Overall, this suggests that tool-first architectures can narrow the performance gap between SLMs and frontier LLMs for many biomedical discovery tasks. While larger models retain advantages in complex multi-turn orchestration and evidence retrieval, smaller models remain competitive in single-turn workflows and provide attractive deployment characteristics for privacy-sensitive, cost-constrained, and enterprise environments.

## 4. Conclusion

This study evaluated the trade-offs between Small Language Models (SLMs) and Large Language Models (LLMs) when used as orchestrators within a biomedical tool-first agent. By holding the workflow, tool ecosystem, prompts, and benchmark constant, the evaluation isolated the impact of the orchestration model on accuracy, efficiency, response quality, and hallucination robustness. The results demonstrated that all evaluated models were capable of effectively orchestrating biomedical workflows, with relatively small differences observed in single-turn tasks and human-evaluated response quality. In particular, Gemma 4 (26B) achieved performance comparable to frontier LLMs across several evaluation dimensions, showing that once scientific knowledge is externalized into tools, databases, and enterprise retrieval systems, SLMs become a realistic option for privacy-preserving, cost-effective scientific agents.

The findings also highlight the effectiveness of the tool-first paradigm for scientific applications. High groundedness scores across all models indicate that responses remained strongly tied to retrieved evidence, reducing dependence on parametric knowledge and mitigating hallucinations. As a result, the primary differences between models were observed in evidence selection, context management, and multi-step workflow execution rather than factual grounding itself. While frontier LLMs retained advantages in complex multi-turn interactions, particularly in evidence validity and workflow management, these gains were achieved at substantially higher computational cost.

More broadly, our findings suggest that future scientific agents should be evaluated not only by the capability of the underlying language model, but also by the quality of the tools they can access and orchestrate. By integrating specialized computational resources within a tool-first architecture, biomedical agents can provide analyses that are more transparent, reproducible, and verifiable than language-model-only approaches. We additionally present practical strategies for improving SLM performance, making them viable orchestrators for biomedical tool ecosystems. Together, these results highlight a path toward cost-effective, privacy-aware, and scientifically grounded agentic systems, while providing guidance on when SLMs or LLMs are most appropriate for a given biomedical application.

## Data availability

The single-turn and multi-turn benchmark datasets used in this study are provided in the Supplementary Data.

## Supplementary Information

**Supplementary information 1**: The complete benchmark dataset of 52 single-turn and 52 multi-turn biomedical tasks, the large language model (LLM)-as-judge prompts used to assess groundedness and evidence validity, and representative case studies examining response quality in relation to token consumption and the impact of prompt design on agent performance. These case studies include model responses, tool-use metrics, latency, token consumption, cost, and evaluation scores.

**Supplementary information 2**:

## References

1 Yang T, Xiao Y, Bao Z, Hao J, Peng J. The rise and potential opportunities of large language model agents in bioinformatics and biomedicine. Brief Bioinform 2025; 26. doi:10.1093/BIB/BBAF601.

2 Li M, Zhao Y, Yu B, Song F, Li H, Yu H et al. API-Bank: A Comprehensive Benchmark for Tool-Augmented LLMs. EMNLP 2023 - 2023 Conference on Empirical Methods in Natural Language Processing, Proceedings 2023; : 3102–3116.

3 Inoue Y, Song T, Wang X, Luna A, Fu T. DruGagent: Multi-Agent Large Language Model-Based Reasoning for Drug-Target Interaction Prediction. ArXiv 2025; : arXiv:2408.13378v4.

4 Belcak P, Heinrich G, Diao S, Fu Y, Dong X, Muralidharan S et al. Small Language Models are the Future of Agentic AI. 2025.https://arxiv.org/pdf/2506.02153 (accessed 14 Oct2026).

5 Patil SG, Mao H, Yan F, Ji CC-J, Suresh V, Stoica I et al. The Berkeley Function Calling Leaderboard (BFCL): From Tool Use to Agentic Evaluation of Large Language Models. In: Proceedings of the 42nd International Conference on Machine Learning. PMLR, 2025, pp 48371–48392.

6 Teams G, Abd S El, Aggarwal V, Algayres R, Andreev A, Bachem O et al. Gemma 4 Technical Report. 2026.https://arxiv.org/pdf/2607.02770 (accessed 14 Oct2026).

7 Comanici G, Bieber E, Schaekermann M, Pasupat I, Sachdeva N, Dhillon I et al. Gemini 2.5: Pushing the Frontier with Advanced Reasoning, Multimodality, Long Context, and Next Generation Agentic Capabilities. 2025.https://arxiv.org/pdf/2507.06261 (accessed 14 Oct2026).

8 Introducing Claude Sonnet 5 \ Anthropic. https://www.anthropic.com/news/claude-sonnet-5 (accessed 14 Oct2026).

9 Zheng L, Chiang WL, Sheng Y, Zhuang S, Wu Z, Zhuang Y et al. Judging LLM-as-a-Judge with MT-Bench and Chatbot Arena. Adv Neural Inf Process Syst 2023; 36. doi:10.52202/075280-2020.

10 Liang P, Bommasani R, Lee T, Tsipras D, Soylu D, Yasunaga M et al. Holistic Evaluation of Language Models. Ann N Y Acad Sci 2022; 1525: 140–146.

11 Wang J, Duan Z. Agent AI with LangGraph: A Modular Framework for Enhancing Machine Translation Using Large Language Models. 2024.https://arxiv.org/pdf/2412.03801 (accessed 14 Oct2026).

12 Ziemann M, Kaspi A, El-Osta A. Digital expression explorer 2: a repository of uniformly processed RNA sequencing data. Gigascience 2019; 8. doi:10.1093/GIGASCIENCE/GIZ022.

13 Love MI, Huber W, Anders S. Moderated estimation of fold change and dispersion for RNA-seq data with DESeq2. Genome Biol 2014; 15. doi:10.1186/S13059-014-0550-8.

14 McEntyre J, Lipman D. PubMed: bridging the information gap. CMAJ: Canadian Medical Association Journal 2001; 164: 1317.

15 Priem J, Piwowar H, Orr R. OpenAlex: A fully-open index of scholarly works, authors, venues, institutions, and concepts. 2022.https://arxiv.org/pdf/2205.01833 (accessed 14 Oct2026).

16 Croft D, O’Kelly G, Wu G, Haw R, Gillespie M, Matthews L et al. Reactome: a database of reactions, pathways and biological processes. Nucleic Acids Res 2011; 39. doi:10.1093/NAR/GKQ1018.

17 Kanehisa M, Furumichi M, Sato Y, Matsuura Y, Ishiguro-Watanabe M. KEGG: biological systems database as a model of the real world. Nucleic Acids Res 2025; 53: D672–D677.

18 Aleksander SA, Balhoff JP, Carbon S, Cherry JM, Ebert D, Feuermann M et al. The Gene Ontology knowledgebase in 2026. Nucleic Acids Res 2026; 54: D1779–D1792.

19 Das D, Krishnan SR, Roy A, Bulusu G. A network-based approach reveals novel invasion and Maurer’s clefts-related proteins in Plasmodium falciparum. Mol Omics 2019; 15: 431– 441.

20 Szklarczyk D, Kirsch R, Koutrouli M, Nastou K, Mehryary F, Hachilif R et al. The STRING database in 2023: protein-protein association networks and functional enrichment analyses for any sequenced genome of interest. Nucleic Acids Res 2023; 51: D638–D646.

21 Chandak P, Huang K, Zitnik M. Building a knowledge graph to enable precision medicine. Sci Data 2023; 10. doi:10.1038/S41597-023-01960-3.

22 Buniello A, Suveges D, Cruz-Castillo C, Llinares MB, Cornu H, Lopez I et al. Open Targets Platform: facilitating therapeutic hypotheses building in drug discovery. Nucleic Acids Res 2025; 53: D1467–D1475.

23 Subramanian A, Narayan R, Corsello SM, Peck DD, Natoli TE, Lu X et al. A Next Generation Connectivity Map: L1000 Platform and the First 1,000,000 Profiles. Cell 2017; 171: 1437–1452.e17.

24 Duan Q, Reid SP, Clark NR, Wang Z, Fernandez NF, Rouillard AD et al. L1000CDS2: LINCS L1000 characteristic direction signatures search engine. npj Systems Biology and Applications 2016 2:1 2016; 2: 16015-.

25 Wang Y, Xiao J, Suzek TO, Zhang J, Wang J, Bryant SH. PubChem: a public information system for analyzing bioactivities of small molecules. Nucleic Acids Res 2009; 37. doi:10.1093/NAR/GKP456.

26 Berman HM, Westbrook J, Feng Z, Gilliland G, Bhat TN, Weissig H et al. The Protein Data Bank. Nucleic Acids Res 2000; 28: 235–242.

27 Jumper J, Evans R, Pritzel A, Green T, Figurnov M, Ronneberger O et al. Highly accurate protein structure prediction with AlphaFold. Nature 2021; 596: 583–589.

28 Volkamer A, Kuhn D, Rippmann F, Rarey M. DoGSiteScorer: a web server for automatic binding site prediction, analysis and druggability assessment. Bioinformatics 2012; 28: 2074–2075.

29 Es S, James J, Espinosa-Anke L, Schockaert S. Ragas: Automated Evaluation of Retrieval Augmented Generation. EACL 2024 - 18th Conference of the European Chapter of the Association for Computational Linguistics, Proceedings of System Demonstrations 2023; : 150–158.

30 Packer C, Wooders S, Lin K, Fang V, Patil SG, Stoica I et al. MemGPT: Towards LLMs as Operating Systems. 2023.https://arxiv.org/pdf/2310.08560 (accessed 14 Oct2026).

31 Zhong W, Guo L, Gao Q, Ye H, Wang Y. MemoryBank: Enhancing Large Language Models with Long-Term Memory. Proceedings of the AAAI Conference on Artificial Intelligence 2023; 38: 19724–19731.

32 Liu W, Huang X, Zeng X, hao xinlong, Yu S, Li D, et al. ToolACE: Winning the Points of LLM Function Calling. International Conference on Learning Representations 2025; 2025: 41359–41381.

