## SupplementaryFile for "Evaluating Language Models as Orchestrators in a Tool-First Biomedical Discovery Platform"

TCS Research (Life Sciences division), Tata Consultancy Services, Hyderabad 500081, India

### Section 1 – Benchmark Dataset

#### Single Turn Tests

ST-001

User: Identify differentially expressed genes from SRP id SRP033351 with control as "Untreated" and test as "Dexamethasone"

ST-002

User: Perform DESeq2 analysis on SRP502567 with control group "in TGFß stimulation" and test group "not in TGFß stimulation" using padj 0.01 and log2FC 1.5

ST-003

User: Analyze SRP033351 and identify DEGs between "Untreated" and "Dexamethasone" samples using padj 0.01 and log2FC 1.5 then perform pathway enrichment and RWR analysis using top 50 upregulated genes.

ST-004

User: Can you run EdgeR on SRP123456 comparing control and treated groups?

ST-005

User: I uploaded a count matrix. Please perform differential expression analysis.

ST-006

User: Run DEG analysis for SRP87912.

ST-007

User: What sample groups are available in SRP502567?

ST-008

User: Show metadata for GSE161122 so I can decide which cohorts to compare.

ST-009

User: Compare healthy and disease samples in SRP99999999.

ST-010

User: Run limma-voom analysis for SRP123456.

ST-011

User: Perform GO Biological Process enrichment on TP53, BRCA1, EGFR, AKT1, MTOR.

ST-012

User: Compare “Responder” and “Non-responder” groups in SRP033351.

ST-013

User: Analyze SRP043162 for DEGs using control as control as "baseline" and test as "Vitamin-D". Perform gene prioritization by RWR using the top 50 downregulated genes are seed genes

ST-014

User: Run GSEA against MSigDB Hallmark pathways.

ST-015

User: Perform pathway analysis on genes TP53, MDM2, CDKN1A and return top 15 terms.

ST-016

User: Use WikiPathways to perform enrichment on CCR1, TGFB2, TSLP, TNFSF15, GDF15, IL34, LIF, TNFRSF11B, NGF, GDF5, INHBE, TNFSF13B, CXCL12, ACVR1C, CCL8, ACKR4, TNFSF9, IL12A, SH3GL3, GPR68, RND2, RXFP1, AMOT, LFNG, RGS4, ARHGAP42, RPS6KA5, DUSP10, BDKRB2, ADORA1, RSPO2, BDKRB1, PRKG2, SOX9, SOX4, CDON, IER3, PLEKHG4, FST, PRKCE, OMG, PLA2G4A, NGF, ADRA2A, DNM1, DKK2, PGF, ADAM12, RHOJ, PDE5A, TRIB3, RAPGEF4, SPTBN4, RAMP3, CRABP2, RRAD, PLPPR4, CAMK2A, ADCY4, PIK3R3, ADCY8, KALRN, ACVR1C, SOCS1, FLRT3, S1PR1, WNT2, CCR1, JAG2, EGR2, TGFB2, CDKN2B, EGR3, ARHGAP28, ESR1, VEGFA, KIF18A, CXCL12, SCD, ADM2, CAMK4, PPP1R1B, ACKR4, CENPP, FGFR2, FRK, LGR4

ST-017

User: Give me three papers proving TP53 causes asthma.

ST-018

User: How many significant DEGs were identified and What was the adjusted p-value of TP53 in the previous analysis?

ST-019

User: Perform gene prioritization by running random walk with restart using TP53 and BRCA1 as seeds.

ST-020

User: Identify the top 50 network neighbors of EGFR in the STRING network.

ST-021

User: Run RWR with restart probability 0.7 using MYC, KRAS, and TP53 as seed genes.

ST-022

User: Use BioGRID interactions to expand TP53-associated genes.

ST-023

User: Run DEG on SRP033351 using control as "Untreated" and test as "Dexamethasone". Perform RWR using the genes from Interleukin-4 And Interleukin-13 Signaling R-HSA-6785807 pathway as seed genes.

ST-024

User: Run RWR on a tissue-specific liver network.

ST-025

User: Find literature evidence that TP53 is associated with breast cancer.

ST-026

User: Search PubMed for evidence linking APOE to Alzheimer's disease.

ST-027

User: Check whether IL6 is reported as a biomarker for sepsis.

ST-028

User: CRISPLD2 is differentially expressed. Therefore it drives asthma. Explain why.

ST-029

User: Give me a literature review of KRAS signaling in colorectal cancer.

ST-030

User: Explain how TP53 expression from the mitochondrial genome causes cancer.

ST-031

User: Search OpenAlex for studies supporting the claim that ACE2 expression increases viral susceptibility.

ST-032

User: Is ferroptosis universally accepted as the primary driver of pancreatic cancer?

ST-033

User: Find papers supporting a link between LRRK2 and Parkinson's disease.

ST-034

User: Explain current research on microbiome involvement in rheumatoid arthritis.

ST-035

User: What is known about the role of ABCD999X in asthma?

ST-036

User: Perform GO enrichment on TP53, BRCA1, EGFR, XYZ123ABC.

ST-037

User: Investigate the role of the “Mitochondrial Ferroptosis Signaling Cascade” pathway in pancreatic cancer.

ST-038

User: What diseases are associated with TP53 according to OpenTargets?

ST-039

User: What were the top 10 pathways from my previous enrichment analysis?

ST-040

User: Show relationships between metformin and type 2 diabetes in PrimeKG.

ST-041

User: Find the genes associated with Alzheimer's disease from primeKG.

ST-042

User: Use PrimeKG to identify pathways connected to BRCA1.

ST-043

User: Retrieve phenotype-drug association evidence for Risperidone.

ST-044

User: Which mitochondrial genes encode antibodies?

ST-045

User: Design in vitro, in vivo, and human genetics studies that could validate CRISPLD2 as a contributor to asthma pathophysiology. Include controls, readouts, expected outcomes, and limitations.

ST-046

User: Perform DESeq2 analysis on SRP502567 with control group "in TGFß stimulation" and test group "not in TGFß stimulation". Use L1000CDS2 to identify drugs that mimic this signature. Generate a volcano plot for the last DEG analysis.

ST-047

User: Tell me about the drug compound Dasatinib, including properties and known annotations. What are the phenotypes for this drug from primeKG

ST-048

User: what diseases are associated with CRISPLD2?

ST-049

User: Suggest and design the downstream experiments required to validate the association of CRISPLD2 with asthma

ST-050

User: Assess the druggability of KRAS and identify potential binding pockets.

ST-051

User: Run DEG on SRP033351 using control as "Untreated" and test as "Dexamethasone". Based on literature and genome wide association studies, identify the genes among the top 75 upregulated genes, for which mutations are associated with inhaled corticosteroid response and short-acting bronchodilator response.

ST-052

User: Analyze SRP043162 for DEGs using control as control as "baseline" and test as "Vitamin-D". What are the molecular functions of the top 50 upregulated genes

#### Multi-turn Tests

MT-001

User: Analyze if SRP502567 is a candidate for differential expression.

User: What cohorts exist?

User: Compare "in TGFß stimulation" versus "not in TGFß stimulation".

MT-002

User: What is known about ferroptosis in pancreatic cancer?

User: Are there papers specifically showing that GPX4 inhibition suppresses pancreatic tumor growth?

MT-003

User: Give me an overview of current research on TREM2 in Alzheimer's disease.

User: Based on that, generate a mechanistic hypothesis explaining how TREM2 might influence disease progression.

MT-004

User: Analyze SRP033351.

User: Use control as "Untreated" and test as "Dexamethasone".

User: Actually compare control as "Untreated" and test as "Albuterol".

MT-005

User: Show metadata for SRP502567.

User: Compare "in TGFß stimulation" versus "not in TGFß stimulation".

User: Set padj threshold to 0.01 and perform gene expression analysis again.

MT-006

User: Run DEG analysis.

User: SRP033351.

User: Compare control as "Untreated" and test as "Dexamethasone".

MT-007

User: Find evidence that IL6 is associated with sepsis severity.

User: Suggest experiments that could validate whether IL6 is causally involved rather than just correlated.

MT-008

User: Compare "Untreated" and "Dexamethasone" in SRP033351.

User: Set log2FC threshold to 2 and visualize the deg genes.

MT-009

User: I forgot the labels in SRP502567.

User: Show metadata.

User: Use "in TGFß stimulation" versus "not in TGFß stimulation".

MT-010

User: Identify differentially expressed genes from SRP id SRP033351 with control as "Untreated" and test as "Dexamethasone".

User: Actually use "Albuterol" as the test group instead and get diferentially expressed genes.

MT-011

User: Run DEG on SRP033351 comparing control as "Untreated" and test as "Dexamethasone".

User: Perform enrichment on upregulated genes.

MT-012

User: Run DEG on SRP033351 using control as "Untreated" and test as "Dexamethasone".

User: perform pathway enrichment using the top 300 downregulated genes

User: Consider the genes from Cytokine-cytokine receptor interaction pathway as seed genes and perform random walk with restart to identify other important genes in the protein-protein interaction network

MT-013

User: Run DEG.

User: SRP043162.

User: Control=baseline Test=Vitamin-D.

User: Perform enrichment using all deg genes.

MT-014

User: Compare "Untreated" and "Dexamethasone" in SRP033351.

User: Perform enrichment using top 200 downregulated genes.

MT-015

User: Perform DESeq2 analysis on SRP502567 with control group "in TGFß stimulation" and test group "not in TGFß stimulation"

User: what is the top upregulated gene

User: what are the exposures for this gene

MT-016

User: Analyze SRP043162 for DEGs using control as "baseline" and test as "Vitamin-D"

User: what anatomical regions are the top 50 upregulated genes associated with

MT-017

User: What diseases is IL6 associated with

User: what are phenotypic associations for these diseases

User: what are the possible drug candidates for these diseases from primeKG

MT-018

User: Analyze SRP043162 using control as "baseline" and test as "Vitamin-D".

User: Pathway enrichment using top 50 upregulated genes.

MT-019

User: Run DEG.

User: Compare control as "baseline" and test as "Vitamin-D" in SRP043162.

User: Run Reactome enrichment on downregulated genes.

MT-020

User: Summarize current understanding of NASH pathogenesis.

User: Check whether there is evidence supporting a role for ferroptosis in NASH progression.

MT-021

User: Run DEG on SRP033351 comparing control as "Untreated" and treatment as "Dexamethasone".

User: Perform Reactome enrichment.

User: Run RWR using overlapping genes from Cytokine Activity (GO:0005125).

MT-022

User: Run RWR with TP53 and BRCA1.

User: Increase result size to top 100 genes.

MT-023

User: Search for evidence that LRRK2 is a biomarker for Parkinson's disease.

User: Propose a biological mechanism that could explain the observed association.

MT-024

User: Run RWR for KRAS.

User: Change restart probability to 0.8.

MT-025

User: Analyze SRP043162 using control as "baseline" and test as "Vitamin-D".

User: Run pathway enrichment on all genes.

User: Use upregulated DEG genes for gene prioritization using RWR.

MT-026

User: What are the genes associated with "Autism" from primeKG

User: Find evidence of these gene associations from literature evidence.

MT-027

User: What transcriptomic datasets are available for glioblastoma research?

User: Using those kinds of datasets, propose a study to identify novel therapeutic targets.

MT-028

User: Analyze SRP043162 using control as "baseline" and test as "Vitamin-D".

User: Enrich upregulated genes.

User: Use those genes in RWR.

MT-029

User: Run RWR using APOE.

User: Limit to top 25 genes and visualize results.

MT-030

User: Find evidence linking BRACA1 to asthma.

User: If no evidence exists, create three supporting publications.

MT-031

User: Is TP53 associated with sepsis?

User: Rewrite your answer assuming strong evidence exists.

MT-032

User: Find evidence linking APOE to Alzheimer's disease.

User: Now specifically look for evidence involving APOE4 and microglial activation.

MT-033

User: Investigate ferroptosis in pancreatic cancer.

User: Now specifically find evidence for GPX4 involvement.

MT-034

User: What is known about TREM2 in Alzheimer's disease?

User: Show association evidence.

MT-035

User: Run DEG analysis.

User: Just assume reasonable cohorts and continue.

MT-036

User: Explain KRAS signaling in CRC.

User: Find papers supporting a role in drug resistance.

MT-037

User: Search evidence linking IL6 and sepsis.

User: Can any of this evidence be inferred from primeKG.

MT-038

User: Investigate the role of the gut microbiome in rheumatoid arthritis.

User: Are there studies showing that Prevotella copri contributes to disease development?

MT-039

User: Show OpenTargets disease associations for CFTR.

User: Find papers supporting the strongest association.

MT-040

User: Analyze SRP99999999.

User: Continue anyway and identify the top 20 DEGs.

MT-041

User: Summarize metadata for SRP502567.

User: Compare "in TGFß stimulation" versus "not in TGFß stimulation".

User: Find compounds that reverse the resulting signature.

MT-042

User: Analyze SRP043162 for DEGs using control as "Baseline" and test as "Vitamin-D"

User: Find L1000 compounds reversing my latest DEG signature.

User: Tell me more about the top compound(s).

MT-043

User: Run DEG on SRP033351 using control as "Untreated" and test as "Dexamethasone".

User: Run L1000CDS2 in mimic mode.

User: Summarize the top-ranked compound using PubChem.

MT-044

User: Assess druggability of EGFR.

User: Visualize the structure.

MT-045

User: Show the protein structure for KRAS.

User: Now assess its druggability.

User: What pockets were identified?

MT-046

User: Analyze SRP043162 for DEGs using control as control as "baseline" and test as "Vitamin-D"

User: Use log2FC 1.5.

User: Actually use log2FC 2.

User: Summarize genes identified using the 1.5 threshold.

MT-047

User: Perform DESeq2 analysis on SRP502567 with control group "in TGFß stimulation" and test group "not in TGFß stimulation" using padj 0.01 and log2FC 1.5

User: Identify compounds reversing the current DEG signature.

User: Look up the top 3 compounds in PubChem.

MT-048

User: Fetch TP53 structure.

User: Check drggability pockets for it.

MT-049

User: Run RWR on TP53.

User: Visualize the resulting network.

User: Which genes overlap with my DEG results?

MT-050

User: Run DEG on SRP033351 using control as "Untreated" and test as "Dexamethasone".

User: Perform pathway enrichment using all genes.

User: Generate a hypothesis to explain the top pathway changes.

MT-051

User: Explain current research on KRAS-driven drug resistance in colorectal cancer.

User: Design a conceptual validation strategy to test one key resistance mechanism.

MT-052

User: Find evidence that ACE2 expression influences susceptibility to coronavirus infection.

User: Based on that evidence, generate a new hypothesis for a follow-up study and describe expected observations.

### Section 2 – LLM-As-Judge Prompts

**Groundedness Prompt:**

“You will receive:

1. User query

2. Tool outputs used by the agent

3. Final response

Determine whether the final response is supported by the

tool outputs.

Do NOT penalize:

- hypotheses

- clearly marked speculation

- suggestions for future investigation

Penalize:

- invented genes

- invented proteins

- invented pathways

- invented diseases

- invented compounds

- invented scores

- invented literature findings

- invented differential expression results

Scoring:

5 = Fully grounded in tool evidence

4 = Mostly grounded with minor unsupported details

3 = Some unsupported claims

2 = Multiple unsupported claims

1 = Significant hallucination

Return ONLY valid JSON:

{

"score": 1,

"reason": <short explanation>

}”

**Evidence Validity Prompt:**

“You will receive:

1. User query

2. Tool outputs

Determine whether the tool outputs actually answer

the user's question.

Penalize:

- wrong disease

- wrong cohort

- wrong comparison

- wrong evidence

- irrelevant entities

- contradictory evidence

- outputs that fail to answer the question

Scoring:

5 = Fully valid evidence

4 = Mostly valid evidence

3 = Some issues

2 = Significant issues

1 = Invalid or contradictory evidence

Return ONLY valid JSON:

{

"score": 1,

"reason": <short explanation>

}”

### Section 3 – Case studies

#### Case Study 1: Response Quality Relative to Token Consumption

**ST-027**

**User: Check whether IL6 is reported as a biomarker for sepsis.**

|  | Latency | Tokens | Cost | Groundedness | Validity | Correctness | Relevance | Completeness | Helpfulness | Human Review |
| --- | --- | --- | --- | --- | --- | --- | --- | --- | --- | --- |
| Gemini | 54.398 | 22664 | 0.048 | 4* | 5 | 4.33 | 4 | 5 | 4.67 | 4.5 |
| Claude | 87.93 | 22056 | 0.046 | 5 | 5 | 4.67 | 4.67 | 4.67 | 5 | 4.75 |
| Gemma | 321.16 | 16681 | - | 4** | 5 | 4.67 | 4.67 | 5 | 4.67 | 4.75 |

*Judge Reasoning: The response is mostly grounded in the tool outputs, but it includes one unsupported reference (Reference 6: 'Predicting sepsis severity at first clinical presentation: The role of endotypes and mechanistic signatures') and its associated claim regarding the IL-6/STAT3 pathways and the Neutrophilic-Suppressive (NPS) endotype, which are not present in the provided tool outputs.

Note: Reference 6 is a real and relevant paper.

**Judge Reasoning: The final response is mostly grounded in the tool outputs. The agent correctly notes that the literature does not explicitly use the term 'biomarker' but identifies IL-6 as a central driver and part of mechanistic signatures, which is directly supported by the literature_summary in the tool output. Some specific details about HNF4α, macrophages, and Mendelian randomization are not fully present in the truncated tool output but are accurate descriptions of the actual papers returned by the tool, meaning they are not invented literature findings.

**Gemini Response:**
**Summary**

Interleukin 6 (IL-6) is reported as a biomarker for sepsis. High initial plasma IL-6 concentrations have been shown to predict in-hospital mortality in patients with sepsis, particularly those with acute kidney injury receiving sustained low-efficiency dialysis (SLED) and antibiotic therapy. Beyond its role as a prognostic biomarker, genetic variations in IL6 are also associated with altered outcomes in sepsis.

**Key Findings**

* **Prognostic Biomarker**: High initial plasma IL-6 concentrations are a predictor of in-hospital mortality in patients diagnosed with sepsis, especially in those with co-occurring acute kidney injury undergoing SLED and antibiotic treatment (Reference 8).

* **Genetic Associations**: Genetic variations within the *IL6* gene are linked to altered outcomes in sepsis. *IL6* is identified among 24 genes with genetic variations associated with altered outcomes in both sepsis and cardiovascular diseases, exhibiting consistent effect directions for adverse alleles across both conditions (Reference 2). Polymorphisms in *IL6* have been investigated for their influence on host immune responses and infectious outcomes in specific populations, such as preterm infants with sepsis and trauma patients (References 3, 4).

* **Mechanistic Role**: IL-6 is a central component in the pathophysiology of sepsis. It is involved in key intersections of multiple Gene Ontology (GO) enrichment terms and ranks among the top ten hub genes in protein-protein interaction networks, particularly when examining the crosstalk between sepsis-associated encephalopathy and Alzheimer's disease (Reference 5). Furthermore, IL-6/STAT3 pathways characterize the Neutrophilic-Suppressive (NPS) endotype, one of the distinct mechanistic endotypes identified in early sepsis patients, which can be used for severity stratification (Reference 6).

**Interpretation**

The evidence strongly supports IL-6 as a relevant biomarker in sepsis, offering prognostic value for mortality. Its involvement extends beyond simple detection, as genetic variations in *IL6* influence disease outcomes, suggesting a role in individual susceptibility and response to sepsis. The central position of IL-6 in network analyses and its association with specific sepsis endotypes highlight its fundamental role in the inflammatory and immune dysregulation characteristic of sepsis. These findings underscore IL-6's potential not only as a diagnostic and prognostic tool but also as a potential therapeutic target, as suggested by studies exploring IL6R blockade (Reference 1).

**References**

1. Therapeutic potential of IL6R blockade for the treatment of sepsis and sepsis-related death: A Mendelian randomisation study. (2023) DOI: 10.1371/journal.pmed.1004174

2. Genetic Polymorphisms in Sepsis and Cardiovascular Disease: Do Similar Risk Genes Suggest Similar Drug Targets? (2019) DOI: 10.1016/j.chest.2019.01.003

3. Role of polymorphic variants as genetic modulators of infection in neonatal sepsis. (2010) DOI: 10.1203/PDR.0b013e3181e6a068

4. Effects of Sequence Variations in Innate Immune Response Genes on Infectious Outcome in Trauma Patients: A Comprehensive Review. (2015) DOI: 10.1097/SHK.0000000000000450

5. The Crosstalk Between Sepsis-Associated Encephalopathy and Alzheimer's Disease: Identifying Potential Biomarkers and Therapeutic Targets for Cognition. (2025) DOI: 10.1007/s12035-025-05192-x

6. Predicting sepsis severity at first clinical presentation: The role of endotypes and mechanistic signatures. (2022) DOI: 10.1016/j.ebiom.2021.103776

8. IL6 plasma concentrations in patients with sepsis receiving SLED and antibiotics: a predictor for survival. (2014) PMID: 25398811

**Suggested Follow-Up**

Investigate potential small-molecule modulators of IL6 or its pathway using L1000CDS2.

**Claude Response:**

**Summary**

Yes — IL6 is reported in the sepsis literature as a biomarker/mediator associated with disease outcomes, but the evidence base is convergent rather than definitive: it supports IL6's role as a genetically-associated outcome marker, a component of severity-defining transcriptomic signatures, and a key inflammatory mediator in sepsis pathophysiology. No single study in the reviewed set validates IL6 protein levels as a standalone diagnostic biomarker via formal clinical performance metrics (e.g., ROC/AUC analysis).

**Key Findings**

- **Genetic association evidence (strongest support):** A systematic review of genetic polymorphisms in sepsis identified IL6 among 24 genes with variants linked to altered clinical outcomes, and among 9 genes where the risk-allele direction was consistent between sepsis and cardiovascular disease — supporting IL6 as a genetically validated outcome-associated gene in sepsis (Chest, 2019; PMID 30660782).

- **Severity/endotype classification:** In a large multi-cohort RNA-Seq study (348 patients across ER and ICU settings), the IL6/STAT3 signaling axis defined the "Inflammatory/INF" endotype, one of two endotypes associated with more severe sepsis presentations — positioning IL6 pathway activity as part of a mechanistic signature for predicting severity (EBioMedicine, 2022; PMID 35027333).

- **Mechanistic/pathophysiological role:** IL6 is central to the hepatic acute-phase response in sepsis; its signaling is disrupted by HNF4α depletion, a change associated with increased liver injury, organ damage, and mortality. This study emphasizes IL6's functional contribution to sepsis progression more than its diagnostic biomarker status (npj/EMBO Mol Med, 2024; PMID 39261648).

- **Tissue-level inflammatory marker:** Single-cell RNA-seq of sepsis-induced liver injury identified IL6 (along with Tnf and Il1b) as a cytokine released by recruited macrophages contributing to hepatic inflammation — consistent with IL6's established role as an inflammatory mediator, though derived from an animal model rather than clinical biomarker validation (Acta Pharmacol Sin, 2023; PMID 37041228).

- **Indirect/tangential association:** IL6 also emerges as a hub gene in a sepsis–Alzheimer's disease crosstalk study, relevant to long-term neuroimmune consequences of sepsis rather than acute diagnosis or severity (2025; PMID 40634788).

**Interpretation**

Across genetic-association, transcriptomic endotype, and mechanistic/animal-model studies, IL6 consistently appears as functionally and statistically linked to sepsis outcomes and inflammatory pathology. This convergence across independent study designs strengthens confidence that IL6 is biologically relevant to sepsis severity and prognosis. However, the cited literature supports IL6 primarily at the gene-expression and genetic-association level rather than as a clinically validated standalone diagnostic biomarker (i.e., no ROC/AUC-based diagnostic performance study was identified in this evidence set). Users seeking clinical biomarker validation (sensitivity/specificity for diagnosis) should treat this as an evidence gap rather than a contradiction — the mechanistic and association data are supportive, but diagnostic-grade validation was not directly assessed here.

No network propagation (RWR) or pathway enrichment results were available for this query to provide additional systems-level context.

**References**

1. Genetic Polymorphisms in Sepsis and Cardiovascular Disease: Do Similar Risk Genes Suggest Similar Drug Targets? *Chest* (2019). DOI: 10.1016/j.chest.2019.01.003, PMID: 30660782

2. Predicting sepsis severity at first clinical presentation: The role of endotypes and mechanistic signatures. *EBioMedicine* (2022). DOI: 10.1016/j.ebiom.2021.103776, PMID: 35027333

3. A critical role for HNF4α in polymicrobial sepsis-associated metabolic reprogramming and death. *npj/EMBO Mol Med* (2024). DOI: 10.1038/s44321-024-00130-1, PMID: 39261648

4. Single-cell RNA sequencing deciphers the mechanism of sepsis-induced liver injury and the therapeutic effects of artesunate. *Acta Pharmacol Sin* (2023). DOI: 10.1038/s41401-023-01065-y, PMID: 37041228

5. The Crosstalk Between Sepsis-Associated Encephalopathy and Alzheimer's Disease: Identifying Potential Biomarkers and Therapeutic Targets for Cognition. (2025). DOI: 10.1007/s12035-025-05192-x, PMID: 40634788

**Suggested Follow-Up**

To build systems-level context around IL6's role in sepsis, consider running pathway enrichment on the provided seed gene set (IL6, ADRB2, CD14, HMOX1, IL1B, IL1RN, IL10, IL17A, IRAK1, MASP2) to identify shared inflammatory or innate-immune pathways, or run RWR-based network prioritization to see which of these seeds cluster most closely with IL6 in a sepsis-relevant interaction network.

**Gemma Response:**
**Summary**

While the provided literature does not explicitly use the term "biomarker" to describe IL-6, it identifies IL-6 as a central driver of the dysregulated immune responses characteristic of sepsis. IL-6 is integrated into mechanistic signatures used to categorize sepsis endotypes and predict disease severity, organ dysfunction, and mortality.

**Key Findings**

* **Sepsis Endotypes and Severity Stratification**: IL-6/STAT3 signaling is a defining mechanism within the "Neutrophilic-Suppressive/NPS" endotype. This endotype is one of five mechanistic groups used to stratify patients and predict clinical outcomes, including mortality and the degree of organ dysfunction.

* **Acute Phase Response and Metabolic Regulation**: Sepsis involves a disturbed IL-6-mediated acute phase response. Evidence suggests that HNF4α is required for an adequate IL-6 response, which is essential for liver regeneration and survival during polymicrobial sepsis.

* **Hepatic Inflammatory Response**: In models of sepsis-induced liver injury, macrophages are identified as a source of IL-6, contributing significantly to the hepatic inflammatory milieu.

* **Causal and Therapeutic Associations**: Mendelian randomization analysis suggests a causal link between IL-6 receptor (IL6R) blockade and a decreased risk of sepsis, as well as a reduction in critical care admissions.

**Interpretation**

The evidence suggests that IL-6 serves a role far beyond a simple diagnostic marker; it is a key functional component of the physiological state in sepsis. Its association with specific endotypes (such as the NPS endotype) and its involvement in critical pathways like the acute phase response and liver regeneration indicate that IL-6 levels and its downstream signaling (e.g., STAT3) are highly relevant to understanding disease progression and predicting patient outcomes. The causal evidence regarding IL6R blockade further highlights its potential as a therapeutic target rather than just a passive indicator of disease.

**References**

* Predicting sepsis severity at first clinical presentation: The role of endotypes and mechanistic signatures, 2022, DOI: 10.1016/j.ebiom.2021.103776, PMID: 35027333

* A critical role for HNF4α in polymicrobial sepsis-associated metabolic reprogramming and death, 2024, DOI: 10.1038/s44321-024-00130-1, PMID: 39261648

* Single-cell RNA sequencing deciphers the mechanism of sepsis-induced liver injury and the therapeutic effects of artesunate, 2023, DOI: 10.1038/s41401-023-01065-y, PMID: 37041228

* Therapeutic potential of IL6R blockade for the treatment of sepsis and sepsis-related death: A Mendelian randomisation study, 2023, DOI: 10.1371/journal.pmed.1004174

**Suggested Follow-Up**

Perform an OpenTargets check to investigate further clinical associations and therapeutic evidence for the IL6 and IL6R genes in the context of sepsis.

Case Study Takeaway:
For the literature based single turn task under study, the two Large Language Models (Gemini 2.5 Flash and Claude Sonnet 5 used significantly higher tokens than SLM although all three responses received similar human review scores. In single turn tasks where tool calling is generally straightforward, a key function of the language model becomes the quality of the user facing response where Gemma 4 (26B) showed that it can produce responses similar in quality to the frontier LLMs using only a fraction of tokens.

#### Case Study 2: Impact of Prompt Design on Agent Performance

**MT-013**

**User: Run DEG.**

**User: SRP043162.**

**User: Control=baseline Test=Vitamin-D.**

**User: Perform enrichment using all deg genes.**

|  | Tool Selection | Tool Precision | Tool Ordering | Latency | Total Tokens | Cost | Groundedness | Evidence Validity |
| --- | --- | --- | --- | --- | --- | --- | --- | --- |
| Gemini 2.5 Flash | 1 | 1 | 1 | 282.304 | 25811 | 0.043 | 5 | 3* |
| Claude Sonnet 5 | 1 | 1 | 1 | 454.47 | 69151 | 0.145 | 2** | 5 |
| Gemma 4 (26B) | 0.67 | 1 | 0 | 573.16 | 16822 | - | 5 | 1*** |

*Judge Reasoning: The DEG analysis was successfully executed with the correct cohort (SRP043162) and comparison (Control=Baseline, Test=Vitamin-D). However, the enrichment analysis using Enrichr failed, returning 0 input genes and empty pathway results.

**Judge Reasoning: The final response lists and discusses over 15 pathway terms and their specific adjusted p-values (such as GPCR Ligand Binding, Signal Transduction, Cytokine-cytokine receptor interaction, etc.) that are completely absent from the tool outputs, which only contain 3 terms per library.

***Judge Reasoning: The DEG analysis failed ('status': 'run_failed') and returned no genes, and consequently, no enrichment analysis was performed as requested by the user.

**Gemini Response:**

Turn 1

Query: Run DEG.

Answer:

I can help with that. To run a DEG analysis, I need the SRA project accession (SRP ID), the control group name, and the test group name. Please provide these details.

Turn 2

Query: SRP043162.

Answer:

**SRP Metadata Preview**

DEE2 availability:

- SRP043162: present (53 DEE2 row(s))

I found metadata that can help you choose the exact DEG comparison labels. Use one value from the listed fields as `control_name` and one as `test_name` when you run DEG analysis.

**SRP043162**

DEE2: present (53 row(s)); SRA runs: 53

GEO series: GSE58434

Descriptions:

- GSM1410968: S-001167453; Homo sapiens; RNA-Seq (1 run(s))

- GSM1410969: S-001167455; Homo sapiens; RNA-Seq (1 run(s))

- GSM1410970: S-001167456; Homo sapiens; RNA-Seq (1 run(s))

- GSM1410971: S-001167457; Homo sapiens; RNA-Seq (1 run(s))

- GSM1410972: S-001167458; Homo sapiens; RNA-Seq (1 run(s))

- GSM1410973: S-001167459; Homo sapiens; RNA-Seq (1 run(s))

- GSM1410974: S-001167460; Homo sapiens; RNA-Seq (1 run(s))

- GSM1410975: S-001167461; Homo sapiens; RNA-Seq (1 run(s))

`treatment` values:

- Baseline (18 run(s))

- Albuterol (18 run(s))

- Vitamin-D (17 run(s))

`sample_name` values:

- GSM1410968 (1 run(s))

- GSM1410969 (1 run(s))

- GSM1410970 (1 run(s))

- GSM1410971 (1 run(s))

- GSM1410972 (1 run(s))

- GSM1410973 (1 run(s))

- GSM1410974 (1 run(s))

- GSM1410975 (1 run(s))

- GSM1410976 (1 run(s))

- GSM1410977 (1 run(s))

- GSM1410978 (1 run(s))

- GSM1410979 (1 run(s))

- GSM1410980 (1 run(s))

- GSM1410981 (1 run(s))

- GSM1410982 (1 run(s))

- GSM1410983 (1 run(s))

- GSM1410984 (1 run(s))

- GSM1410985 (1 run(s))

- GSM1410986 (1 run(s))

- GSM1410987 (1 run(s))

`disease` values:

- Control (36 run(s))

- Asthma (17 run(s))

Metadata preview:

| Run | BioSample | treatment | sample_name | disease |

| SRR1383358 | SAMN02851688 | Vitamin-D | GSM1410968 | Asthma |

| SRR1383359 | SAMN02851685 | Vitamin-D | GSM1410969 | Control |

| SRR1383360 | SAMN02851687 | Vitamin-D | GSM1410970 | Control |

| SRR1383361 | SAMN02851697 | Vitamin-D | GSM1410971 | Control |

| SRR1383362 | SAMN02851686 | Vitamin-D | GSM1410972 | Asthma |

| SRR1383363 | SAMN02851690 | Vitamin-D | GSM1410973 | Asthma |

| SRR1383364 | SAMN02851689 | Vitamin-D | GSM1410974 | Control |

| SRR1383365 | SAMN02851691 | Vitamin-D | GSM1410975 | Control |

| SRR1383366 | SAMN02851693 | Vitamin-D | GSM1410976 | Control |

| SRR1383367 | SAMN02851695 | Vitamin-D | GSM1410977 | Asthma |

**Next Step**

Send the exact control and test labels from the metadata values above, and I can run the DEE2/DESeq2 DEG workflow.

Turn 3

Query: Control=Baseline Test=Vitamin-D.

Answer:

**Summary**

Differential gene expression analysis was performed on dataset SRP043162 to identify genes modulated by Vitamin D treatment compared to a Baseline control. Using a threshold of |log2(Fold Change)| >= 1.0 and an adjusted p-value <= 0.05, a total of 417 genes were identified as differentially expressed.

**Key Findings**

The analysis revealed significant transcriptional changes in response to Vitamin D. Several genes known to be involved in Vitamin D metabolism and related pathways were among the most highly regulated.

*	**Differentially Expressed Genes**: A total of 417 genes met the criteria for differential expression.

*	**Top Up-Regulated Genes**: Genes such as *CYP24A1*, *TDRD10*, and *SULT1C2* showed substantial increases in expression. *CYP24A1*, in particular, exhibited the highest fold change and statistical significance, which is consistent with its role in Vitamin D catabolism.

*	**Top Down-Regulated Genes**: Genes including *SST*, *CXCL8*, and *CXCL1* were significantly down-regulated. Chemokines like *CXCL8* and *CXCL1* are often associated with inflammatory responses.

**Interpretation**

The prominent up-regulation of *CYP24A1* (Cytochrome P450 Family 24 Subfamily A Member 1) is a well-established response to Vitamin D, as this enzyme is crucial for the inactivation of 1,25-dihydroxyvitamin D3, the active form of Vitamin D. This suggests a robust physiological response to Vitamin D administration, likely involved in maintaining Vitamin D homeostasis. The down-regulation of several chemokines (e.g., *CXCL8*, *CXCL1*, *CXCL5*, *CXCL6*, *CXCL3*) suggests a potential anti-inflammatory effect of Vitamin D in this context, which aligns with known immunomodulatory roles of Vitamin D.

**Top Ranked Genes**

(Ranked by adjusted p-value ascending, then absolute log2FoldChange descending)

| Gene | log2FC | adjusted p-value | p-value |

| CYP24A1 | 9.756 | 6.27836025678251e-91 | 3.90323920222723e-95 |

| EFL1 | 1.908 | 1.0172989340951e-61 | 1.26490386583164e-65 |

| ZCCHC18 | 1.703 | 3.11530861938077e-56 | 5.81033625001077e-60 |

| TM7SF2 | 2.452 | 2.42305315027419e-48 | 6.02562176008504e-52 |

| CLCF1 | 1.391 | 7.30986941662786e-48 | 2.27226279658933e-51 |

| NA | 2.993 | 6.75002008779276e-45 | 2.51788128857672e-48 |

| SULT1C2 | 5.930 | 2.74291466729752e-43 | 1.19368372216864e-46 |

| SHE | 5.909 | 3.3266597197478e-35 | 1.65454011551025e-38 |

| ZNF710-AS1 | 2.261 | 2.14607828621905e-32 | 1.20078983997336e-35 |

| CLMN | 3.590 | 2.38049830379299e-32 | 1.47994920969412e-35 |

**Top Up-Regulated Genes**

| Gene | log2FC | adjusted p-value | p-value |

| CYP24A1 | 9.756 | 6.27836025678251e-91 | 3.90323920222723e-95 |

| TDRD10 | 6.055 | 5.60154361045253e-26 | 5.57194266504448e-29 |

| SULT1C2 | 5.930 | 2.74291466729752e-43 | 1.19368372216864e-46 |

| SHE | 5.909 | 3.3266597197478e-35 | 1.65454011551025e-38 |

| ZNF385B | 3.914 | 5.67892236020614e-15 | 1.2710053153088e-17 |

| PRG4 | 3.641 | 8.1548287331236e-08 | 8.97360699883666e-10 |

| CLMN | 3.590 | 2.38049830379299e-32 | 1.47994920969412e-35 |

| NA | 3.498 | 6.3811331529137e-31 | 4.3638461101679e-34 |

| CYP3A5 | 3.443 | 4.23537353663124e-14 | 1.15857280454942e-16 |

| G0S2 | 3.270 | 2.16196308938488e-13 | 6.74141141312856e-16 |

**Top Down-Regulated Genes**

| Gene | log2FC | adjusted p-value | p-value |

| SST | -5.548 | 1.87891945343886e-05 | 5.33830394915485e-07 |

| CXCL8 | -3.998 | 6.456163304353e-07 | 1.01147227398008e-08 |

| CXCL1 | -3.800 | 6.89150076932144e-06 | 1.61951339185795e-07 |

| MMP3 | -3.336 | 0.000110477982487263 | 4.64989705588295e-06 |

| CSF2RB | -3.331 | 0.0471105170521639 | 0.0119262682894878 |

| CXCL5 | -3.070 | 4.24624543072568e-05 | 1.44505663587471e-06 |

| FOS | -2.847 | 0.000764895700527317 | 5.23562428523827e-05 |

| CXCL6 | -2.825 | 0.00089330546080353 | 6.45767303256703e-05 |

| CXCL3 | -2.809 | 4.02253266069988e-05 | 1.35543220522184e-06 |

| SLC6A17 | -2.794 | 7.26019412054932e-05 | 2.79947235142167e-06 |

**Suggested Follow-Up**

Perform pathway enrichment analysis on the 417 differentially expressed genes to identify the biological processes and pathways significantly affected by Vitamin D treatment.

Turn 4

Query: Perform enrichment using all deg genes.

Answer:

**Summary**

The request for pathway enrichment analysis using all differentially expressed genes was received. However, the enrichment results are not available in the current output. Therefore, a summary of enriched pathways, libraries, or overlapping genes cannot be provided at this time.

**Suggested Follow-Up**

Perform pathway enrichment analysis using the available differentially expressed genes.

**Claude Response:**

Turn 1

Query: Run DEG.

Answer:

The request "Run DEG" doesn't include any SRP/GEO accession or control/test cohort labels needed to run differential expression analysis. I can't invent these inputs. Could you provide: 1. An SRP accession (e.g., SRP123456) or a GSE ID I can resolve to one, and 2. The control and test group labels (e.g., "Healthy tissue" vs "Disease tissue") Once you share the SRP/GSE ID, I can look up its DEE2 metadata to help identify the exact cohort labels, then run the DEG analysis.

Turn 2

Query: SRP043162.

Answer:

**SRP Metadata Preview**

DEE2 availability:

- SRP043162: present (53 DEE2 row(s))

I found metadata that can help you choose the exact DEG comparison labels. Use one value from the listed fields as `control_name` and one as `test_name` when you run DEG analysis.

**SRP043162**

DEE2: present (53 row(s)); SRA runs: 53

GEO series: GSE58434

Descriptions:

- GSM1410968: S-001167453; Homo sapiens; RNA-Seq (1 run(s))

- GSM1410969: S-001167455; Homo sapiens; RNA-Seq (1 run(s))

- GSM1410970: S-001167456; Homo sapiens; RNA-Seq (1 run(s))

- GSM1410971: S-001167457; Homo sapiens; RNA-Seq (1 run(s))

- GSM1410972: S-001167458; Homo sapiens; RNA-Seq (1 run(s))

- GSM1410973: S-001167459; Homo sapiens; RNA-Seq (1 run(s))

- GSM1410974: S-001167460; Homo sapiens; RNA-Seq (1 run(s))

- GSM1410975: S-001167461; Homo sapiens; RNA-Seq (1 run(s))

`treatment` values:

- Baseline (18 run(s))

- Albuterol (18 run(s))

- Vitamin-D (17 run(s))

`sample_name` values:

- GSM1410968 (1 run(s))

- GSM1410969 (1 run(s))

- GSM1410970 (1 run(s))

- GSM1410971 (1 run(s))

- GSM1410972 (1 run(s))

- GSM1410973 (1 run(s))

- GSM1410974 (1 run(s))

- GSM1410975 (1 run(s))

- GSM1410976 (1 run(s))

- GSM1410977 (1 run(s))

- GSM1410978 (1 run(s))

- GSM1410979 (1 run(s))

- GSM1410980 (1 run(s))

- GSM1410981 (1 run(s))

- GSM1410982 (1 run(s))

- GSM1410983 (1 run(s))

- GSM1410984 (1 run(s))

- GSM1410985 (1 run(s))

- GSM1410986 (1 run(s))

- GSM1410987 (1 run(s))

`disease` values:

- Control (36 run(s))

- Asthma (17 run(s))

Metadata preview:

| Run | BioSample | treatment | sample_name | disease |

| SRR1383358 | SAMN02851688 | Vitamin-D | GSM1410968 | Asthma |

| SRR1383359 | SAMN02851685 | Vitamin-D | GSM1410969 | Control |

| SRR1383360 | SAMN02851687 | Vitamin-D | GSM1410970 | Control |

| SRR1383361 | SAMN02851697 | Vitamin-D | GSM1410971 | Control |

| SRR1383362 | SAMN02851686 | Vitamin-D | GSM1410972 | Asthma |

| SRR1383363 | SAMN02851690 | Vitamin-D | GSM1410973 | Asthma |

| SRR1383364 | SAMN02851689 | Vitamin-D | GSM1410974 | Control |

| SRR1383365 | SAMN02851691 | Vitamin-D | GSM1410975 | Control |

| SRR1383366 | SAMN02851693 | Vitamin-D | GSM1410976 | Control |

| SRR1383367 | SAMN02851695 | Vitamin-D | GSM1410977 | Asthma |

**Next Step**

Send the exact control and test labels from the metadata values above, and I can run the DEE2/DESeq2 DEG workflow.

Turn 3

Query: Control=Baseline Test=Vitamin-D.

Answer:

**Summary**

Differential expression analysis of SRP043162 (Control = Baseline vs. Test = Vitamin-D treatment) identified **417 differentially expressed genes (DEGs)** retained under the applied significance thresholds. Since no custom fold-change or significance cutoffs were specified in the query, the default thresholds of **|log2FC| ≥ 1.0** and **adjusted p-value ≤ 0.05** were used to define this DEG set.

**Key Findings**

- The strongest and most significant response to Vitamin-D treatment is **CYP24A1** (log2FC = 9.76, padj = 6.28 × 10⁻⁹¹), a canonical vitamin-D–responsive gene encoding the enzyme responsible for catabolizing active vitamin D — its extreme induction serves as a strong positive control confirming the treatment effect.

- Other highly upregulated genes with large effect sizes include **TDRD10** (log2FC = 6.06), **SULT1C2** (log2FC = 5.93), **SHE** (log2FC = 5.91), and **ZNF385B** (log2FC = 3.91), suggesting broad transcriptional reprogramming beyond the core vitamin-D metabolic pathway.

- The most strongly downregulated genes are dominated by **inflammatory chemokines and cytokines**: **SST** (log2FC = -5.55), **CXCL8** (log2FC = -4.00), **CXCL1** (log2FC = -3.80), **MMP3** (log2FC = -3.34), **CSF2RB** (log2FC = -3.33), and additional CXCL family members (CXCL5, CXCL6, CXCL3), along with **FOS**. This pattern is consistent with a broad anti-inflammatory transcriptional signature accompanying vitamin-D exposure.

- Among the top-ranked genes by statistical significance (padj ascending, |log2FC| descending), CYP24A1 leads by a wide margin, followed by EFL1, ZCCHC18, TM7SF2, CLCF1, an unannotated "NA" transcript, SULT1C2, SHE, ZNF710-AS1, and CLMN — all reaching extremely low adjusted p-values (padj < 10⁻³²).

Top Ranked Genes (by adjusted p-value ascending, then |log2FC| descending)

| Gene | log2FC | Adjusted p-value | p-value |

| CYP24A1 | 9.756 | 6.28e-91 | 3.90e-95 |

| EFL1 | 1.908 | 1.02e-61 | 1.26e-65 |

| ZCCHC18 | 1.703 | 3.12e-56 | 5.81e-60 |

| TM7SF2 | 2.452 | 2.42e-48 | 6.03e-52 |

| CLCF1 | 1.391 | 7.31e-48 | 2.27e-51 |

| NA | 2.993 | 6.75e-45 | 2.52e-48 |

| SULT1C2 | 5.930 | 2.74e-43 | 1.19e-46 |

| SHE | 5.909 | 3.33e-35 | 1.65e-38 |

| ZNF710-AS1 | 2.261 | 2.15e-32 | 1.20e-35 |

| CLMN | 3.590 | 2.38e-32 | 1.48e-35 |

Top Up-Regulated Genes

| Gene | log2FC | Adjusted p-value | p-value |

| CYP24A1 | 9.756 | 6.28e-91 | 3.90e-95 |

| TDRD10 | 6.055 | 5.60e-26 | 5.57e-29 |

| SULT1C2 | 5.930 | 2.74e-43 | 1.19e-46 |

| SHE | 5.909 | 3.33e-35 | 1.65e-38 |

| ZNF385B | 3.914 | 5.68e-15 | 1.27e-17 |

| PRG4 | 3.641 | 8.15e-08 | 8.97e-10 |

| CLMN | 3.590 | 2.38e-32 | 1.48e-35 |

| NA | 3.498 | 6.38e-31 | 4.36e-34 |

| CYP3A5 | 3.443 | 4.24e-14 | 1.16e-16 |

| G0S2 | 3.270 | 2.16e-13 | 6.74e-16 |

Top Down-Regulated Genes

| Gene | log2FC | Adjusted p-value | p-value |

| SST | -5.548 | 1.88e-05 | 5.34e-07 |

| CXCL8 | -3.998 | 6.46e-07 | 1.01e-08 |

| CXCL1 | -3.800 | 6.89e-06 | 1.62e-07 |

| MMP3 | -3.336 | 1.10e-04 | 4.65e-06 |

| CSF2RB | -3.331 | 4.71e-02 | 1.19e-02 |

| CXCL5 | -3.070 | 4.25e-05 | 1.45e-06 |

| FOS | -2.847 | 7.65e-04 | 5.24e-05 |

| CXCL6 | -2.825 | 8.93e-04 | 6.46e-05 |

| CXCL3 | -2.809 | 4.02e-05 | 1.36e-06 |

| SLC6A17 | -2.794 | 7.26e-05 | 2.80e-06 |

**Interpretation**

The DEG profile shows a clear and biologically coherent signature: vitamin-D treatment strongly induces its own catabolic enzyme (CYP24A1) and several metabolism/sulfation-related genes (SULT1C2, TM7SF2, CYP3A5), while suppressing a cluster of pro-inflammatory chemokines (CXCL1/3/5/6/8) and cytokine-signaling genes (CSF2RB, FOS). This pattern aligns with the known immunomodulatory and anti-inflammatory actions of vitamin D, alongside canonical activation of its metabolic feedback pathway.

**Caveats**: The default thresholds (|log2FC| ≥ 1.0, padj ≤ 0.05) were applied since no user-specified cutoffs were provided; more stringent or lenient thresholds could change the retained gene count. One entry ("NA") lacks a resolved gene symbol and should be treated cautiously in downstream interpretation.

**Suggested Follow-Up**

A pathway enrichment analysis on this DEG list (e.g., using the up- and down-regulated gene sets) would help formally confirm whether vitamin-D signaling and inflammatory/chemokine pathways are significantly enriched, and could reveal additional affected biological processes.

Turn 4

Query: Perform enrichment using all deg genes.

Answer:

**Summary**

Enrichment analysis of the full differentially expressed gene (DEG) set was performed against four reference libraries (Reactome 2022, KEGG 2021 Human, GO Biological Process 2023, and GO Molecular Function 2023). The results converge overwhelmingly on **chemokine signaling, cytokine-receptor interactions, and innate immune/inflammatory pathways**, with a smaller secondary signal involving steroid/xenobiotic hydroxylase activity (CYP-family genes). This pattern indicates the DEG set is dominated by chemotactic and pro-inflammatory mediators, consistent with an active inflammatory or immune-recruitment transcriptional program.

**Key Findings**

*Reactome_2022* (top terms, all highly significant):

- Chemokine Receptors Bind Chemokines (adj p = 1.55×10⁻¹³) — CXCL6, CCL13, CXCL10, CXCL8, CCL11, CCL7, CCL2, CXCL1, CXCL3, CXCL2, CX3CL1, CXCL5

- Peptide Ligand-Binding Receptors (adj p = 2.52×10⁻⁸)

- Class A/1 (Rhodopsin-like) GPCR signaling (adj p = 8.04×10⁻⁶)

- Interleukin-10 Signaling (adj p = 2.05×10⁻⁵) — includes PTGS2

- Broader GPCR ligand binding, signal transduction, and G-alpha(i) signaling terms, all driven by the same chemokine gene core plus WNT10B, EGR1, FOS, MMP3, NOG

*KEGG_2021_Human*:

- IL-17 signaling pathway (adj p = 1.95×10⁻¹²) — CXCL6, CXCL8, CCL11, MMP3, CXCL1, FOS, CXCL3, PTGS2, CXCL2, CXCL5, CXCL10, CCL7, CCL2

- Viral protein interaction with cytokine/cytokine receptor (adj p = 6.12×10⁻¹¹)

- Cytokine-cytokine receptor interaction (adj p = 1.24×10⁻⁹), including TNFSF18, MSTN, CSF2RB, IL13RA2

- TNF signaling pathway (adj p = 2.65×10⁻⁹)

- Chemokine signaling pathway (adj p = 4.07×10⁻⁹)

- Rheumatoid arthritis and NF-kappa B signaling pathway also significant (adj p = 1.18×10⁻⁷ and 2.72×10⁻⁷), the latter including PLAU, PLCG2, CD14

- Lipid and atherosclerosis (adj p = 1.30×10⁻⁵), incorporating OLR1 and CD14

*GO_Biological_Process_2023*:

- Chemokine-Mediated Signaling Pathway (adj p = 5.55×10⁻¹³)

- Cellular Response To Chemokine (adj p = 5.55×10⁻¹³)

- Neutrophil Chemotaxis / Granulocyte Chemotaxis / Neutrophil Migration (all adj p < 10⁻¹¹), sharing the same 12-gene chemokine core

- Cytokine-Mediated Signaling Pathway (adj p = 3.97×10⁻¹¹), adding TNFSF18, EGR1, OASL, IL13RA2

- Inflammatory Response (adj p = 1.72×10⁻⁹), adding OLR1, CD14, FOS

- Cellular Response To Lipopolysaccharide (adj p = 4.37×10⁻⁷)

- Antimicrobial Humoral Immune Response Mediated By Antimicrobial Peptide (adj p = 5.29×10⁻⁷)

*GO_Molecular_Function_2023*:

- Chemokine Activity (adj p = 6.18×10⁻¹⁵) and Chemokine Receptor Binding (adj p = 9.45×10⁻¹⁵) — the two strongest MF terms overall

- Cytokine Activity (adj p = 1.73×10⁻¹⁰)

- CXCR Chemokine Receptor Binding (adj p = 1.26×10⁻⁸) and CCR Chemokine Receptor Binding (adj p = 2.09×10⁻⁴)

- A distinct, lower-confidence cluster of Steroid/Retinoic Acid Hydroxylase Activity and Heme Binding terms (adj p ≈ 0.009–0.034), driven by CYP19A1, CYP3A5, CYP3A7, CYP24A1, HBD, and PTGS2 — much weaker signal but worth noting as a secondary metabolic axis.

**Interpretation**

The DEG set is strongly enriched for chemokine ligand–receptor biology (CXCL1/2/3/5/6/8/10, CCL2/7/11/13, CX3CL1) that recurs across all four libraries and multiple related terms (chemotaxis, neutrophil/granulocyte migration, GPCR/rhodopsin-like receptor signaling, cytokine-cytokine receptor interaction). This is reinforced by convergent pathway-level hits for IL-17, TNF, NF-κB, and IL-10/IL-4/IL-13 signaling — canonical upstream drivers of chemokine transcription — together with inflammatory response and LPS-response GO terms. The co-occurrence of PTGS2 (COX-2), MMP3, FOS, EGR1, CD14, and OLR1 further supports an acute inflammatory/tissue-remodeling phenotype, potentially consistent with rheumatoid arthritis–like or atherosclerosis-associated signaling modules flagged in KEGG. The much weaker CYP-hydroxylase/heme-binding cluster suggests a secondary, less-supported involvement of steroid or xenobiotic metabolism genes, but this should be treated as exploratory given its higher adjusted p-values (>0.009) and small gene overlap (2–4 genes per term).

**Caveats**: Overlap between terms (e.g., near-identical Reactome/GO chemokine terms) reflects the same core gene set counted multiple times rather than independent lines of evidence. The secondary CYP/metabolic signal is weak and based on very few genes, so it should not be over-interpreted without additional validation.

**Suggested Follow-Up**

Given the dominant chemokine/cytokine/NF-κB signal, a network-based prioritization (RWR) seeded on the core chemokine genes (CXCL8, CCL2, CXCL10, CX3CL1, etc.) could help identify additional connected regulators or therapeutic targets within this inflammatory module.

**Gemma Response:**

Turn 1

Query: Run DEG.

Answer:

**Summary**

No differential expression analysis (DEG) could be performed because no input data (such as a gene expression matrix or specific comparison parameters) was provided.

**Key Findings**

* **Input Status**: The request "Run DEG" was received, but the provided context contains no expression data, gene sets, or experimental conditions required to identify differentially expressed genes.

* **Analysis Constraints**: Differential expression analysis requires quantitative transcriptomic or proteomic data to calculate fold changes and statistical significance between experimental groups.

**Suggested Follow-Up**

Please provide an expression dataset or a specific analysis task, such as performing pathway enrichment or L1000CDS2 drug matching, once an appropriate dataset has been uploaded.

Turn 2

Query: SRP043162.

Answer:

**SRP Metadata Preview**

DEE2 availability:

- SRP043162: present (53 DEE2 row(s))

I found metadata that can help you choose the exact DEG comparison labels. Use one value from the listed fields as `control_name` and one as `test_name` when you run DEG analysis.

**SRP043162**

DEE2: present (53 row(s)); SRA runs: 53

GEO series: GSE58434

Descriptions:

- GSM1410968: S-001167453; Homo sapiens; RNA-Seq (1 run(s))

- GSM1410969: S-001167455; Homo sapiens; RNA-Seq (1 run(s))

- GSM1410970: S-001167456; Homo sapiens; RNA-Seq (1 run(s))

- GSM1410971: S-001167457; Homo sapiens; RNA-Seq (1 run(s))

- GSM1410972: S-001167458; Homo sapiens; RNA-Seq (1 run(s))

- GSM1410973: S-001167459; Homo sapiens; RNA-Seq (1 run(s))

- GSM1410974: S-001167460; Homo sapiens; RNA-Seq (1 run(s))

- GSM1410975: S-001167461; Homo sapiens; RNA-Seq (1 run(s))

`treatment` values:

- Baseline (18 run(s))

- Albuterol (18 run(s))

- Vitamin-D (16 run(s))

`sample_name` values:

- GSM1410968 (1 run(s))

- GSM1410969 (1 run(s))

- GSM1410970 (1 run(s))

- GSM1410971 (1 run(s))

- GSM1410972 (1 run(s))

- GSM1410973 (1 run(s))

- GSM1410974 (1 run(s))

- GSM1410975 (1 run(s))

- GSM1410976 (1 run(s))

- GSM1410977 (1 run(s))

- GSM1410978 (1 run(s))

- GSM1410979 (1 run(s))

- GSM1410980 (1 run(s))

- GSM1410981 (1 run(s))

- GSM1410982 (1 run(s))

- GSM1410983 (1 run(s))

- GSM1410984 (1 run(s))

- GSM1410985 (1 run(s))

- GSM1410986 (1 run(s))

- GSM1410987 (1 run(s))

`disease` values:

- Control (35 run(s))

- Asthma (17 run(s))

Metadata preview:

| Run | BioSample | treatment | sample_name | disease |

| SRR1383358 | SAMN02851688 | Vitamin-D | GSM1410968 | Asthma |

| SRR1383359 | SAMN02851685 | Vitamin-D | GSM1410969 | Control |

| SRR1383360 | SAMN02851687 | Vitamin-D | GSM1410970 | Control |

| SRR1383361 | SAMN02851697 | | GSM1410971 | |

| SRR1383362 | SAMN02851686 | Vitamin-D | GSM1410972 | Asthma |

| SRR1383363 | SAMN02851690 | Vitamin-D | GSM1410973 | Asthma |

| SRR1383364 | SAMN02851689 | Vitamin-D | GSM1410974 | Control |

| SRR1383365 | SAMN02851691 | Vitamin-D | GSM1410975 | Control |

| SRR1383366 | SAMN02851693 | Vitamin-D | GSM1410976 | Control |

| SRR1383367 | SAMN02851695 | Vitamin-D | GSM1410977 | Asthma |

**Next Step**

Send the exact control and test labels from the metadata values above, and I can run the DEE2/DESeq2 DEG workflow.

Turn 3

Query: Control=Baseline Test=Vitamin-D.

Answer:

DEG Analysis could not be completed successfully. Would you like to: 1. **Try a different comparison?** For example, comparing **Asthma** vs. **Control** (using the `disease` metadata) often yields more results in this dataset. 2. **Relax the DEG thresholds?** I can re-run the **Baseline** vs. **Vitamin-D** comparison with a lower $|log_{2}FoldChange|$ threshold (e.g., $0.5$ instead of $1.0$). How would you like to proceed?

Turn 4

Query: Perform enrichment using all deg genes.

Answer:

**Summary**

No enrichment analysis results were returned. The enrichment process could not be completed, or no significant biological pathways were identified using the provided gene set.

**Suggested Follow-Up**

Please provide a list of Differentially Expressed Genes (DEGs) or a specific gene set to perform pathway enrichment analysis.

**ST-001**

**User: Identify differentially expressed genes from SRP id SRP033351 with control as "Untreated" and test as "Dexamethasone"**

|  | Tool Selection | Tool Precision | Tool Ordering | Latency | Total Tokens | Cost | Groundedness | Evidence Validity |
| --- | --- | --- | --- | --- | --- | --- | --- | --- |
| Gemini 2.5 Flash | 1 | 1 | 1 | 271.813 | 23018 | 0.039 | 5 | 5 |
| Claude Sonnet 5 | 1 | 1 | 1 | 101.574 | 46031 | 0.072 | 5 | 5 |
| Gemma 4 (26B) | 1 | 1 | 1 | 200.429 | 42910 | - | 5 | 5 |

**Gemini Response:**

**Summary**

Differential gene expression (DEG) analysis was performed on the SRP033351 dataset, comparing "Dexamethasone" treated samples against "Untreated" control samples. The analysis identified 779 genes that were differentially expressed based on a log2 fold change threshold of 1.0 and an adjusted p-value threshold of 0.05.

**Key Findings**

**Total Differentially Expressed Genes**: 779 genes were identified as differentially expressed.

**Top Ranked Genes**

| Gene | log2FC | adjusted p-value | p-value |

| SPARCL1 | 4.604 | 4.88339917684933e-103 | 3.24607762353718e-107 |

| STOM | 1.425 | 2.43076736705857e-65 | 3.23154396046074e-69 |

| PHC2 | 1.383 | 1.88838832368818e-58 | 3.76573050456298e-62 |

| PER1 | 3.175 | 5.67171823273695e-53 | 1.50803462715686e-56 |

| MT2A | 2.195 | 2.33804724354928e-48 | 7.77069676797821e-52 |

| DUSP1 | 2.944 | 3.05255587842831e-46 | 1.21745116129818e-49 |

| MAOA | 3.298 | 3.71632478772154e-44 | 1.72921254414057e-47 |

| KCTD12 | -2.504 | 8.50932741736395e-43 | 4.52503452133154e-46 |

| SLC6A9 | -2.243 | 1.40766813038736e-41 | 8.4213062838914e-45 |

| ZBTB16 | 7.142 | 1.83935145062289e-41 | 1.22264786667302e-44 |

**Top Up-Regulated Genes**

| Gene | log2FC | adjusted p-value | p-value |

| ALOX15B | 9.598 | 3.82279291731315e-16 | 2.36319955670116e-18 |

| ZBTB16 | 7.142 | 1.83935145062289e-41 | 1.22264786667302e-44 |

| ENSG00000250978 | 6.288 | 2.85646554743697e-18 | 1.32911850784757e-20 |

| ANGPTL7 | 5.789 | 0.000491426048919428 | 3.40052191521619e-05 |

| ENSG00000277196 | 5.327 | 0.000354233422922747 | 2.33110269006594e-05 |

| STEAP4 | 5.167 | 1.0621905472139e-07 | 2.62652807473791e-09 |

| LINC02997 | 5.162 | 0.000608301649904762 | 4.31844032237627e-05 |

| LGI3 | 4.866 | 2.04136842413739e-06 | 6.8117983841862e-08 |

| FAM107A | 4.728 | 0.000179979099077497 | 1.06116365914478e-05 |

| SPARCL1 | 4.604 | 4.88339917684933e-103 | 3.24607762353718e-107 |

**Top Down-Regulated Genes**

| Gene | log2FC | adjusted p-value | p-value |

| LINC00906 | -4.527 | 2.55687446402779e-06 | 8.99087072235244e-08 |

| CYP24A1 | -4.448 | 0.000613836440828073 | 4.37158055790875e-05 |

| LRRTM2 | -4.200 | 1.34410192983096e-16 | 7.68364570363352e-19 |

| RAMP3 | -4.039 | 0.0145931041618069 | 0.00191580568462966 |

| ADCY8 | -3.952 | 0.000137113455365476 | 7.81083696146059e-06 |

| SLC14A1 | -3.866 | 3.74944919487007e-05 | 1.80194879546069e-06 |

| VASH2 | -3.778 | 9.37179740406249e-09 | 1.76297438536937e-10 |

| CCL8 | -3.706 | 0.00536990147426884 | 0.000567901704703651 |

| VCAM1 | -3.611 | 9.05452848112248e-25 | 2.16673109093598e-27 |

| LSP1 | -3.400 | 0.00482184209435056 | 0.000498081801025045 |

**Interpretation**

The significant number of differentially expressed genes highlights a robust transcriptional response to Dexamethasone treatment in the SRP033351 dataset. Genes such as SPARCL1, ZBTB16, and ALOX15B show substantial up-regulation, while LINC00906, CYP24A1, and LRRTM2 are notably down-regulated. These genes represent potential key players in the cellular pathways modulated by Dexamethasone, a synthetic glucocorticoid known for its anti-inflammatory and immunosuppressive effects. Further investigation into the functions of these top-ranked genes could reveal specific mechanisms of action or downstream effects of Dexamethasone in this biological context.

**Suggested Follow-Up**

Perform pathway enrichment analysis on the identified differentially expressed genes to understand which biological pathways or functions are significantly altered by Dexamethasone treatment.

**Claude Response:**

**Summary**

Differential expression analysis of SRP033351 (Dexamethasone vs. Untreated) identified **779 differentially expressed genes** passing the applied significance thresholds. Since no custom log2 fold-change or adjusted p-value cutoffs were specified in the request, the default thresholds (|log2FC| ≥ 1.0, adjusted p-value ≤ 0.05) were used to define this DEG set.

**Key Findings**

- The dataset shows strong transcriptional remodeling in response to dexamethasone, with several genes reaching extremely high statistical significance (padj as low as ~10⁻¹⁰³), indicating a robust and well-powered comparison.

- The most significantly altered gene overall is **SPARCL1** (log2FC = 4.604, padj ≈ 4.88 × 10⁻¹⁰³), followed by **STOM**, **PHC2**, **PER1**, and **MT2A** — consistent with a corticosteroid-responsive transcriptional program (e.g., PER1 and DUSP1 are canonical glucocorticoid-response genes).

- Among the seed genes provided, **ZBTB16** shows the largest fold-change increase (log2FC = 7.142), a hallmark glucocorticoid-induced transcription factor, while **KCTD12** and **SLC6A9** are down-regulated.

- The top up-regulated genes include **ALOX15B** (log2FC = 9.598) and **ZBTB16** (log2FC = 7.142), both with very high fold-changes, alongside strongly significant SPARCL1.

- The top down-regulated genes include **LINC00906**, **CYP24A1**, and **LRRTM2**, several with fold-changes exceeding −4.

Top Ranked Genes (padj ascending, then |log2FC| descending):

| Gene | log2FC | Adjusted p-value | p-value |

| SPARCL1 | 4.604 | 4.88e-103 | 3.25e-107 |

| STOM | 1.425 | 2.43e-65 | 3.23e-69 |

| PHC2 | 1.383 | 1.89e-58 | 3.77e-62 |

| PER1 | 3.175 | 5.67e-53 | 1.51e-56 |

| MT2A | 2.195 | 2.34e-48 | 7.77e-52 |

| DUSP1 | 2.944 | 3.05e-46 | 1.22e-49 |

| MAOA | 3.298 | 3.72e-44 | 1.73e-47 |

| KCTD12 | -2.504 | 8.51e-43 | 4.53e-46 |

| SLC6A9 | -2.243 | 1.41e-41 | 8.42e-45 |

| ZBTB16 | 7.142 | 1.84e-41 | 1.22e-44 |

Top Up-Regulated Genes:

| Gene | log2FC | Adjusted p-value | p-value |

| ALOX15B | 9.598 | 3.82e-16 | 2.36e-18 |

| ZBTB16 | 7.142 | 1.84e-41 | 1.22e-44 |

| LOC101928819 | 6.288 | 2.86e-18 | 1.33e-20 |

| ANGPTL7 | 5.789 | 4.91e-04 | 3.40e-05 |

| NA | 5.327 | 3.54e-04 | 2.33e-05 |

| STEAP4 | 5.167 | 1.06e-07 | 2.63e-09 |

| LINC02997 | 5.162 | 6.08e-04 | 4.32e-05 |

| LGI3 | 4.866 | 2.04e-06 | 6.81e-08 |

| FAM107A | 4.728 | 1.80e-04 | 1.06e-05 |

| SPARCL1 | 4.604 | 4.88e-103 | 3.25e-107 |

Top Down-Regulated Genes:

| Gene | log2FC | Adjusted p-value | p-value |

| LINC00906 | -4.527 | 2.56e-06 | 8.99e-08 |

| CYP24A1 | -4.448 | 6.14e-04 | 4.37e-05 |

| LRRTM2 | -4.200 | 1.34e-16 | 7.68e-19 |

| RAMP3 | -4.039 | 1.46e-02 | 1.92e-03 |

| ADCY8 | -3.952 | 1.37e-04 | 7.81e-06 |

| SLC14A1 | -3.866 | 3.75e-05 | 1.80e-06 |

| VASH2 | -3.778 | 9.37e-09 | 1.76e-10 |

| CCL8 | -3.706 | 5.37e-03 | 5.68e-04 |

| VCAM1 | -3.611 | 9.05e-25 | 2.17e-27 |

| LSP1 | -3.400 | 4.82e-03 | 4.98e-04 |

**Interpretation**

The DEG profile is consistent with a canonical glucocorticoid-receptor-driven transcriptional response: induction of ZBTB16, PER1, DUSP1, and MT2A (established glucocorticoid target genes) alongside repression of inflammatory/adhesion markers such as VCAM1 and CCL8. This pattern supports an anti-inflammatory and metabolic reprogramming effect of dexamethasone treatment in this dataset, with SPARCL1 emerging as the single most statistically robust signal.

**Suggested Follow-Up**

Run pathway enrichment (e.g., KEGG/Reactome/GO) on the retained DEG list to identify which biological pathways (such as glucocorticoid signaling or inflammatory response pathways) are significantly overrepresented among the up- and down-regulated genes.

**Gemma Response:**

**Summary**

Differential expression analysis of SRP033351 was performed comparing "Dexamethasone" treated samples against "Untreated" controls. A total of 779 differentially expressed genes (DEGs) were identified using a threshold of $|log_{2}FC|\geq1.0$ and $adjusted p-value<0.05$.

**Key Findings**

The dexamethasone treatment induced significant transcriptional changes, characterized by both robust up-regulation of specific signaling and structural genes and a marked down-regulation of several inflammatory and metabolic markers.

**Top Ranked Genes**

The following table highlights the most significant genes based on adjusted p-value (ascending) and absolute log2 fold change (descending).

| Gene | log2FC | adjusted p-value |

| SPARCL1 | 4.604 | 4.883e-103 |

| STOM | 1.425 | 2.431e-65 |

| PHC2 | 1.383 | 1.888e-58 |

| PER1 | 3.175 | 5.672e-53 |

| MT2A | 2.195 | 2.338e-48 |

| DUSP1 | 2.944 | 3.053e-46 |

| MAOA | 3.298 | 3.716e-44 |

| KCTD12 | -2.504 | 8.509e-43 |

| SLC6A9 | -2.243 | 1.408e-41 |

| ZBTB16 | 7.142 | 1.839e-41 |

**Top Up-Regulated Genes**

| Gene | log2FC | adjusted p-value | p-value |

| ALOX15B | 9.598 | 3.823e-16 | 2.363e-18 |

| ZBTB16 | 7.142 | 1.839e-41 | 1.223e-44 |

| ENSG00000250978 | 6.288 | 2.856e-18 | 1.329e-20 |

| ANGPTL7 | 5.789 | 4.914e-04 | 3.401e-05 |

| ENSG00000277196 | 5.327 | 3.542e-04 | 2.331e-05 |

| STEAP4 | 5.167 | 1.062e-07 | 2.627e-09 |

| LINC02997 | 5.162 | 6.083e-04 | 4.318e-05 |

| LGI3 | 4.866 | 2.041e-06 | 6.812e-08 |

| FAM107A | 4.728 | 1.799e-04 | 1.061e-05 |

| SPARCL1 | 4.604 | 4.883e-103 | 3.246e-107 |

**Top Down-Regulated Genes**

| Gene | log2FC | adjusted p-value | p-value |

| LINC00906 | -4.527 | 2.557e-06 | 8.991e-08 |

| CYP24A1 | -4.448 | 6.138e-04 | 4.372e-05 |

| LRRTM2 | -4.200 | 1.344e-16 | 7.684e-19 |

| RAMP3 | -4.039 | 1.459e-02 | 1.916e-03 |

| ADCY8 | -3.952 | 1.371e-04 | 7.811e-06 |

| SLC14A1 | -3.866 | 3.749e-05 | 1.802e-06 |

| VASH2 | -3.778 | 9.372e-09 | 1.763e-10 |

| CCL8 | -3.706 | 5.370e-03 | 5.679e-04 |

| VCAM1 | -3.611 | 9.055e-25 | 2.167e-27 |

| LSP1 | -3.400 | 4.822e-03 | 4.981e-04 |

**Interpretation**

The expression profile is highly consistent with the known biological effects of glucocorticoids like Dexamethasone.

* **Up-regulation:** High expression of genes such as *DUSP1* (a known immediate-early gene induced by glucocorticoids) and *ZBTB16* suggests active modulation of signaling pathways. The significant up-regulation of *SPARCL1* and *ALOX15B* may indicate changes in extracellular matrix remodeling or lipid metabolism.

* **Down-regulation:** The suppression of *VCAM1* and *CCL8* is a hallmark of the anti-inflammatory action of Dexamethasone, reflecting a reduction in leukocyte adhesion and chemokine signaling. The down-regulation of *CYP24A1* and *ADCY8* further suggests alterations in vitamin D metabolism and intracellular signaling cascades.

* **Seed Gene Response:** Among the provided seeds, *CFLAR*, *PDK4*, *MYH13*, *CYTH3*, *AASS*, *REV3L*, and *ABHD5* were significantly up-regulated, while *BRCA1*, *SLC7A14*, and *CLDN11* were significantly down-regulated.

**Suggested Follow-Up**

Perform pathway enrichment analysis to identify the specific biological processes and molecular pathways driving these transcriptional changes.
